# Sex-specific transcript diversity is regulated by a maternal transcription factor in early *Drosophila* embryos

**DOI:** 10.1101/2021.03.18.436074

**Authors:** Mukulika Ray, Ashley Mae Conard, Jennifer Urban, Joseph Aguilera, Annie Huang, Pranav Mahableshwarkar, Smriti Vaidyanathan, Erica Larschan

## Abstract

Co-transcriptional splicing coordinates the processes of transcription and splicing and is driven by transcription factors (TFs) and diverse RNA-binding proteins (RBPs). Yet the mechanisms by which specific TFs and RBPs function together in context-specific ways to drive precise co-transcriptional splicing at each of thousands of genomic loci remains unknown. Therefore, we have used sex-specific splicing in *Drosophila* as a model to understand how the function of TFs and RBPs is coordinated to transcribe and process specific RNA transcripts at the correct genomic locations. We show widespread sex-specific transcript diversity occurs much earlier than previously thought and present a new pipeline called time2splice to quantify splicing changes over time. We define several mechanisms by which the essential and functionally-conserved CLAMP TF functions with specific RBPs to precisely regulate co-transcriptional splicing: 1) CLAMP links the DNA of gene bodies of sex-specifically spliced genes directly to the RNA of target genes and physically interacts with snRNA and protein components of the splicing machinery; 2) In males, CLAMP regulates the distribution of the highly conserved RBP **M**ale**le**ss (MLE) (RNA Helicase A) to prevent aberrant sex-specific splicing; 3) In females, CLAMP modulates alternative splicing by directly binding to target DNA and RNA and indirectly through regulating the splicing of *sex lethal*, the master regulator of sex determination. Overall, we provide new insight into how TFs function specifically with RBPs to drive alternative splicing.

## Introduction

One of the greatest challenges in modern biology is to understand how transcription is coupled with splicing to drive development and differentiation. Gene expression is a multistep process, initiated when RNA transcripts are synthesized from DNA templates, followed at many genes by **a**lternative **s**plicing (AS), the selective inclusion or exclusion of introns and exons. Immediately after **t**ranscription **f**actors (TFs) initiate transcription, multiple **R**NA **B**inding **P**roteins (RPBs) bind to nascent RNA and remove introns to form the mature mRNA, ready for export and translation^1–3^. Thus, both TFs and RBPs drive transcriptome diversity through regulating transcript levels and isoforms. These processes are likely coupled, as much prior work^2, 4–6^ and recent cryo-EM structures^7^ reveal a close association between TFs and splicing complexes. Furthermore, many RBPs that are part of splicing complexes, regulate both transcription and splicing^8–12^. However, it is not yet understood how TFs and RBPs function together to coordinate transcription and alternative splicing such that specific isoforms are transcribed and processed at the correct genomic locations. Therefore, we have studied the coordination of transcription and alternative splicing during the establishment of sexual dimorphism in *Drosophila melanogaster* as a model to understand this process.

A key to understanding how alternative splicing is established as sexual dimorphism initiates lies in the events that shape the initial few hours of an organism’s existence. During early development, TFs and RBPs deposited by the mother into the embryo shape early embryonic milestones across metazoans^13, 14^. Initially, cell number increases, followed by differentiation into specific cell types, which is driven by transcription and co-transcriptional splicing^3, 15^. However, the mechanisms by which maternally deposited TFs and RBPs function together to drive the process of co-transcriptional splicing in a sex-specific manner remains poorly understood.

The *Drosophila* embryo is an excellent tool to study the role of maternally deposited proteins and RNA in early development as it is easy to perform genetic manipulations to remove maternal factors to define how they regulate splicing and transcription. Also, embryos can be sexed before zygotic genome activation due to our recent application of a meiotic drive system^16^. During *Drosophila* embryogenesis, zygotic genome activation (ZGA) occurs shortly after the first two hours of development. Concurrently, maternal transcripts gradually decrease in abundance, and zygotic transcription increases, a process called the MZT (**M**aternal to **Z**ygotic **T**ransition). ZGA starts approximately 80 min after egg laying and most maternal transcripts are degraded by 180 min after egg laying^17^. Even at these early stages of development, AS generates multiple transcript isoforms resulting in transcript diversity. Although the earliest genes transcribed from the zygotic genome are mainly intron-less, approximately 30% of early zygotic transcripts do have introns^18, 19^. Furthermore, genes involved in sex determination do have introns and use AS to drive male versus female-specific development^20^. Hence, during early embryonic development, AS is important for shaping cell and tissue-specific transcriptomes and essential for sexual differentiation. However, it was not known whether maternally-deposited factors initiate sex-specific AS early in development.

TFs have been hypothesized to facilitate context-specific AS, but little is known about how specific transcription factors mediate this link. Several lines of evidence led us to hypothesize that the maternally-deposited TF CLAMP (**C**hromatin-**l**inked **a**dapter for **M**SL **p**roteins) is a good candidate with which to study the mechanisms by which TFs and RBPs function together to drive sex-specific co-transcriptional AS for several reasons: 1) CLAMP directly binds to DNA at both promoters and intronic sites^21, 22^ and mass spectrometry identified association with 33 RBPs on chromatin, 6 of which regulate AS^23^; 2) CLAMP is bound to approximately equal numbers of intronic regions and promoter regions^24^; 3) Many CLAMP binding sites evolved from intronic polypyrimidine tracts^25^; 4) Maternal CLAMP is essential for viability in males and females^22^. Therefore, we hypothesized that CLAMP is a maternally deposited TF that selectively interacts with specific RBPs to regulate co-transcriptional AS in a context-specific manner.

We combined diverse genomic, genetic, and computational approaches to define new mechanisms that control the context-specificity of co-transcriptional splicing using sex-specific splicing as a model. First, we determined all of the sex-specifically spliced isoforms early during development genome-wide which has never been performed in any species. Next, we identified the following mechanisms: 1) In both males and females, CLAMP acts as a linker between the DNA of gene bodies and the RNA of a subset of its targeted sex-specifically spliced genes; 2) CLAMP sex-specifically interacts with spliceosomal RNAs and RBPs; 3) In males specifically, CLAMP regulates the distribution of the spliceosome component MLE (**M**ale**le**ss) on chromatin to prevent aberrant sex-specific splicing; 4) In females specifically, CLAMP functions upstream of the *sxl* master regulator of sex determination to directly regulate splicing and directly and indirectly regulate the AS of different subsets of downstream *sxl* targets. Thus, we conclude that we have identified new mechanisms by which a TF functions context-specifically with RBPs to regulate alternative splicing that drives a key developmental decision.

## Results

### 1. Sex-specific alternative splicing is present at the earliest stages of Drosophila development

Before we could define the function of TFs and RBPs in regulating sex-specific splicing, it was critical to define when sex-specific splicing begins during development. Therefore, we analyzed RNA-sequencing data that we generated from male and female embryos at two-time points: 0-2 hours (pre-MZT) and 2-4 hours (post-MZT)^16^ (#GSE102922). We were able to produce male or female embryos using a meiotic drive system that produces sperm with either only X or only Y chromosomes^16^. Next, we measured the amount of AS in these samples using a new pipeline that we developed for this analysis and made publicly available called time2Splice (https://github.com/ashleymaeconard/time2splice). Time2Splice implements the commonly used SUPPA2 algorithm^26^ to identify splice variants and provides additional modules to integrate time, sex, and chromatin localization data (**Materials and Methods**) (**Fig S1**). SUPPA2 measures the PSI (**P**ercent **S**pliced **I**n) for each exon and calculates the differential alternative splicing between samples, reported as ΔPSI^26^. Therefore, SUPPA2 is specifically designed to identify alternative splicing events.

From our RNA-seq data, we used Time2Splice to analyze 66,927 exons associated with 17,558 genes and classified the AS events into one of seven classes (diagrammed in **Fig 1A**). We found that 16-18% of the exons are alternatively spliced in early embryos (**Fig1B)** and fall into one of the seven classes (**Fig 1C-D**). Of these seven classes, AF (**A**lternative **F**irst Exon) is the most common type, constituting almost one-fourth of total AS (∼24-26%), and AL (**A**lternative **L**ast Exon) is the least common type (∼3%). The AS transcript distribution across categories was similar between the two time points and the two sexes (**Fig 1B-D**). Next, we asked which type of AS is most affected by depleting CLAMP. The overall distribution of transcripts into the seven AS classes remains mostly unaffected in the absence of maternal CLAMP. However, at the 0-2 Hr (pre-MZT) stage, loss of maternal CLAMP results in a more substantial decrease in **M**utually E**x**clusive **E**xon (**MXE**) splicing in both males and females compared with all of the other types of splicing (**males**: p-value < 3.21e-21; **females**: p-value < 6.26e-87 Chi-squared test) (**Fig 1D**). At the 2-4 Hr/post-MZT stage, only male embryos have a significant percentage of MXE splicing affected in the absence of maternal CLAMP (p-value < 1.95e-137 Chi-squared test) (**Fig 1D**). Therefore, CLAMP regulates AS and has a stronger effect on MXE splicing than other types of splicing.

**Fig 1.**
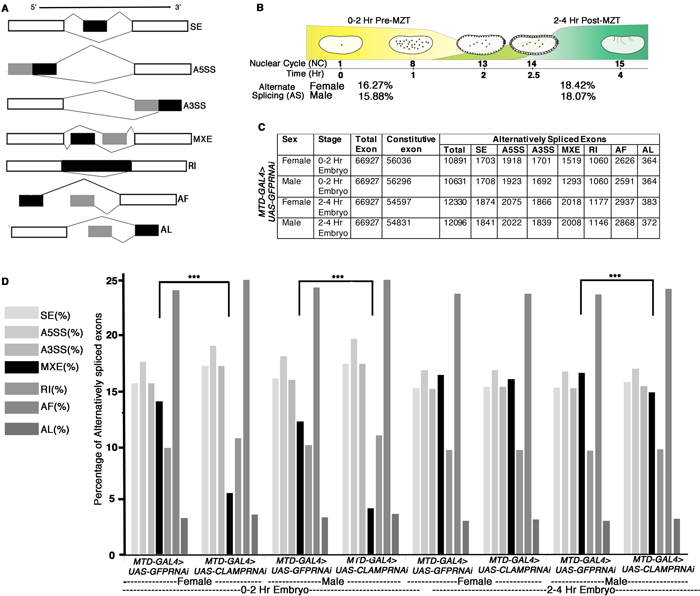
Alternative splicing during early *Drosophila melanogaster* embryonic development. **A.** Schematic diagrams showing seven different types of Alternative splicing (AS). The constitutive exons are depicted as white rectangles, whereas the alternatively spliced exons are in shades of black and grey rectangles **B.** Percentage of genes with alternative splicing in male and female early *Drosophila* embryos at the 0-2 Hr/pre-MZT and 2-4 Hr/post-MZT stages. **C.** Table showing the number of exons in each AS category in control sexed embryos at the 0-2 Hr/pre-MZT and 2-4 Hr/post-MZT stages. **D.** Bar plot showing the distribution of different types of AS at 0-2 Hr/pre-MZT and 2-4 Hr/ post-MZT for female and male embryos in the presence (*MTDGAL4>UAS-GFPRNAi*) and absence (*MTDGAL4>UAS-CLAMPRNAi*) of maternal CLAMP. A Chi-square test was performed to determine if there is a significant difference between the percentage of each type of AS including MXE splicing (black bar) in the presence vs. absence of CLAMP in each class of sample: female and male 0-2 Hr/pre-MZT, and 2-4 Hr/post-MZT embryos. Statistically significant differences (p<0.001 marked by ***) were found between categories connected by solid black lines.

During MXE splicing, one isoform of the transcript retains one of the alternative exons and excludes another exon, which is retained by another isoform (schematic in **Fig 1A**). Interestingly, MXE alternative splicing occurs in many transcripts that encode components of the sex determination pathway^12^. Furthermore, CLAMP has a sex-specific role in dosage compensation^27, 28^. Therefore, we defined sex-specific splicing events in the early embryo for the first time. We identified sex-specific splicing events in 0-2 Hr embryos (pre-MZT) (**Fig S2A, N=92**) and in 2-4 Hr embryos (post-MZT) (**Fig S2B, N=138**) and categorized them as known **s**ex-**s**pecifically **s**pliced (**known SSS**) events. Overall, we determined that sex-specific AS occurs earlier in development than ever shown previously in any species.

### 2. Maternal CLAMP regulates sex-specific alternative splicing in early Drosophila embryos

We hypothesized that CLAMP regulates sex-specific AS in early embryos for the following reasons: 1) CLAMP is a maternally deposited pioneer transcription factor with sex-specific functions that is enriched at intronic regions in addition to promoters^24, 29^; 2) Proteomic data identified a physical association between spliceosome components and CLAMP^23^; and 3) CLAMP binding sites evolved from polypyrimidine tracts that regulate splicing^25^. We tested our hypothesis in early staged and sexed embryos by measuring differences in splicing in RNA-seq data generated from male and female 0-2 Hr/pre-MZT and 2-4 Hr/post-MZT embryos with and without maternal CLAMP^16^. The maternal triple driver GAL4 (*MTD-GAL4*) was used to drive *UAS-CLAMPRNAi[val22]* which strongly reduces maternal CLAMP levels as validated by qPCR and Western blot conducted in parallel with mRNA-seq data collection ^16^.

First, we asked whether CLAMP alters AS and we found 200-400 transcripts at which AS is regulated by CLAMP depending on the time point and sex (**Fig S2C-F** and **Fig 2A, B**). To determine whether CLAMP-dependent AS events are enriched for **s**ex-**s**pecific **s**plicing (SSS) events, we first identified all of the CLAMP dependent AS events in female (**Fig S2C, D**) and in male (**Fig S2E, F**) 0-2 Hr and 2-4 Hr embryos (**Materials and Methods**). We measured alternative splicing using an exon-centric approach to quantify individual splice junctions by measuring PSI (**P**ercent **S**pliced **I**n) for a particular exon using the established SUPPA algorithm within the time2splice pipeline^26^. Exon inclusion is represented as positive PSI, and exclusion events are defined as negative PSI (equation in **Materials and Methods**). By comparing the CLAMP-dependent AS events in females and males, we identified CLAMP-dependent sex-specific splicing events in female and male in 0-2 Hr and 2-4 Hr embryos (**Fig 2A, B and Supplementary Table S1**).

**Fig 2.**
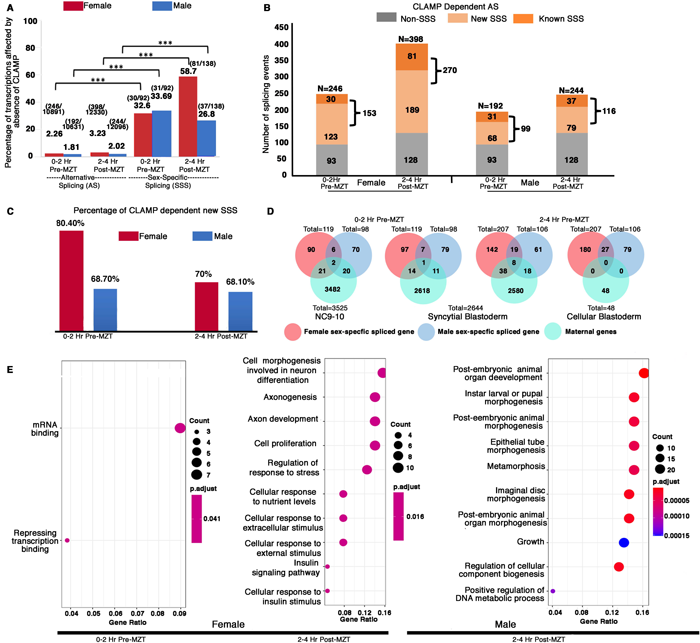
Maternal CLAMP regulates sex-specific alternative splicing during early embryonic development. **A.** Bar graph showing the percentage of transcripts (raw values noted at the top of each bar) compared with total AS events or sex-specific splicing (SSS) events within parentheses listed at the top of each bar: number of splicing events regulated by CLAMP/Total number of splicing events. We quantified transcripts whose splicing is regulated by maternal CLAMP at the 0-2Hr/pre-MZT and 2-4Hr/post-MZT stages in females (red bars) and males (blue bars). A Fisher’s Exact Test was performed with significance shown at p<0.001. **B.** Bar plot showing the total number of splicing events undergoing CLAMP-dependent AS (N) in females and males at 0-2 Hr/pre-MZT and 2-4 Hr/post-MZT embryonic stages. Alternatively, spliced genes are divided into non-sex-specific (grey) and sex-specific (orange shades) sub-categories. CLAMP-dependent female and male sex-specifically spliced (SSS) genes are divided into known (sex-specific in control samples: darker orange) and new (sex-specific only after depleting CLAMP: lighter orange) sub-categories identified from 0-2 Hr/pre-MZT and 2-4 Hr post-MZT /embryos. **C.** Percentage of **new** female (red) and male (blue) CLAMP-dependent sex-specifically spliced genes in 0-2 Hr/pre-MZT and 2-4 Hr/post-MZT embryos that were not identified as different between males and females in control samples. **D.** Female (red) and male (blue) CLAMP-dependent sex-specific spliced genes compared with maternal genes (green, NC9-10 stage, **N=3525**; Syncytial Blastoderm stage, **N=2644**; Cellular Blastoderm stage, **N=48**) at 0-2 Hr/pre-MZT (female, **N=119** and male, **N=98**) and 2-4 Hr/ post-MZT stages (female, **N= 207** and male, **N=106**). **E.** Gene Ontology (GO) results for genes showing CLAMP-dependent female sex-specific splicing in embryos at the 0-2 Hr/pre-MZT stage and for genes exhibiting CLAMP-dependent female and male sex-specific splicing in embryos at the 2-4 Hr/post-MZT stage. The size of the circle increases as the number of genes in that category increases. The color of the circle represents significance (p-value). GO categories for male embryos at the 0-2 Hr/pre-MZT stage are not shown because the gene set is small and therefore no enriched GO categories were identified.

When we measured the percentage of total alternatively spliced and sex-specifically spliced transcripts that are CLAMP-dependent in males and females at both pre-and post-MZT stages, we found that while only 2-3% of total AS exons are CLAMP-dependent, ∼30-60% of sex-specifically spliced exons are CLAMP-dependent (**Fig 2A**). Therefore, the function of CLAMP in AS is highly enriched at sex-specific transcripts. We then divided all CLAMP-dependent AS events into two categories: 1) sex-specifically spliced (SSS) events; and 2) non-sex specifically spliced (non-SSS) events (**Fig 2B**). We then subdivided the CLAMP-dependent SSS events into the following subclasses: 1) **known** SSS events, female-specific and male-specific splicing events at 0-2 Hr and 2-4 Hr embryo stages that are CLAMP dependent (p<0.05) (**Fig 2A-B**); 2) **new** SSS events, splicing events that occur only in the absence of CLAMP and not in control samples (**Fig 2B**), which are aberrant splicing events suppressed by CLAMP. By calculating ι1PSI in these subclasses, we identified widespread CLAMP-dependent sex-specific splicing, especially in female embryos (**Fig 2B**). Interestingly, the majority of CLAMP-dependent SSS events are **new** aberrant SSS events that did not occur in the presence of maternal CLAMP (∼70%) (**Fig 2C**). Furthermore, 85% of genes at which CLAMP regulates sex-specific splicing, it does not regulate transcription as determined by comparing our differentially spliced genes with our published differential gene expression analysis^16^. Therefore, the function of CLAMP in regulating sex-specific splicing largely does not overlap with its regulation of transcription.

To define the magnitude of the effect of CLAMP on splicing, we compared the ι1PSI for known and new SSS events between female and male samples (**Fig S3**). We found that although more splicing events/transcripts show CLAMP-dependent splicing in females (∼150-250) than males (∼100) (**Fig 2B and Supplementary Table S1**), post-MZT, CLAMP-dependent exon inclusion was significantly enriched in male new SSS transcripts compared to their female-specific counterparts (**Fig S3**). Thus, in the absence of CLAMP, new aberrant sex-specifically spliced isoforms are generated suggesting that CLAMP normally inhibits aberrant sex-specific splicing events.

During the first few hours of their development, *Drosophila* embryos have predominantly maternal transcripts. Therefore, we asked whether CLAMP-dependent female and male specifically-spliced genes are maternally deposited or zygotically transcribed. We compared our list of CLAMP-dependent sex-specifically spliced genes with known maternally expressed genes that are consistent across multiple previous studies^30, 31^. We found very low levels of overlap with maternally deposited transcripts (**Fig 2D**) even in the 0-2 Hr embryo stage, consistent with ZGA starting at approximately 80 minutes after egg laying^17^. Therefore, most of the sex-specifically spliced genes we observed are likely to be zygotic transcripts, consistent with the function of CLAMP as a pioneer TF acting in the early embryo^22^.

To understand the classes of genes whose splicing is CLAMP-dependent, we performed Gene Ontology (GO) analysis. Our analysis showed that pre-MZT (0-2 Hrs), female-specifically spliced genes are primarily TFs and factors that regulate splicing (**Fig 2E**). Therefore, in females CLAMP alters the splicing of genes that can regulate the transcription and splicing of other genes that amplify its regulatory role. In contrast, the male specifically-spliced pre-MZT genes are not enriched for any specific biological function or process, likely due to the small number of genes in the gene list. At the post-MZT stage in both sexes, CLAMP regulates the splicing of genes that drive development including organogenesis, morphogenesis, cell proliferation, signaling, and neurogenesis (**Fig 2E**).

In order to validate our genomic splicing analysis from time2splice for individual target genes (**Fig. 2E**), we randomly selected eight genes at which we determined CLAMP regulates splicing using qRT-PCR or RT-PCR (**Fig. S4**). Our RT-PCR results indicate that we are able to validate the function of CLAMP in regulating splicing of genes which we identified genomically with time2splice (**Fig. S4**). We summarized the functions of the validated target genes at which splicing is regulated by CLAMP (**Supplementary Table S2**). *iab4,* one of the target genes that we validated, has known functional links to CLAMP suggesting that we have identified relevant target genes^23, 29, 32^. Furthermore, many of the validated target genes that are sex-specifically spliced by CLAMP are themselves involved in splicing and chromatin regulation including those with known isoforms that specifically regulate alternative splicing such as *fus*, *pep*, *sc35* (**Supplementary Table S2**). In summary, maternal CLAMP functions early in development to prevent the aberrant sex-specific splicing of the majority of sex-specifically spliced zygotic transcripts including many that encode regulators of alternative splicing.

### 3. Zygotic CLAMP regulates sex-specific alternative splicing during Drosophila development

Next, we asked whether in addition to maternal CLAMP, zygotic CLAMP regulates sex-specific splicing. Therefore, we analyzed total RNA-seq data from wild type control and *clamp* null mutant (*clamp*^2^)^27^ third instar larvae (L3) and identified CLAMP dependent sex-specific splicing events (**Supplementary Table S3**). Out of a total of 189 and 211 CLAMP-dependent alternative splicing events in female and male L3 larvae, we identified 139/189 (73.5%) and 161/211 (76.3%) sex-specific splicing events (**Supplementary Table S3, Sheet1, Column H-J**). Because CLAMP regulates transcription in addition to splicing, we compared transcription and splicing target genes to each other and found that 60% of sex-specifically spliced genes are also regulated at the level of transcription in larvae in contrast to 15% of sex-specifically spliced genes in embryos (**Supplementary Table S4, Fig S5A, S5B**). This shows that CLAMP has a dual role in transcription and splicing that differs at different developmental stages.

Zygotic CLAMP is also present in male and female cell lines derived from embryos. Such cell lines have frequently been used as model systems due to the ability to easily obtain sufficient material for low yield genomic assays like ChIP-seq^33^, MNAse-seq^21, 34^ and iCLIP (see below). Furthermore, S2 and Kc cells are embryonically-derived established models for male and female cells, respectively, that differ in their sex chromosome complement^35, 36^ and have been studied for decades in the context of dosage compensation^37–39^. Therefore, we also defined CLAMP-dependent splicing events by performing *clamp* RNAi in Kc and S2 cells. We first quantified all splicing events that differ between control populations of Kc and S2 cells using time2splice (**Supplementary Table S5, Fig S5C)**. Then we identified CLAMP-dependent splicing events in cell lines and found that these events are almost entirely sex-specific: 1) 45/46 CLAMP-dependent splicing events are female sex-specific in Kc cells and 112/113 CLAMP-dependent splicing events are male sex-specific in S2 cells (**Supplementary Table S5, Sheet2, Column F-H**).

Interestingly, almost all CLAMP-dependent spliced genes were regulated by CLAMP at the level of splicing and not transcription and many more genes are regulated at the level of splicing than transcription (**Fig S5D)**. While 100 genes (112 splicing events) show CLAMP-dependent splicing in S2 cells, only 12 genes exhibit CLAMP-dependent differential gene expression. Similarly, in Kc cells, 42 genes (45 splicing events) show CLAMP-dependent splicing and only 18 genes show CLAMP-dependent expression (**Fig S5D, E**). Overall, fewer genes are regulated by CLAMP in cell lines compared with embryos likely because cell lines remain alive in the absence of CLAMP^40^ while embryos depleted for maternal CLAMP do not survive past zygotic genome activation and L3 null larvae do not undergo pupation^29^. In summary, zygotic CLAMP regulates splicing in larvae and embryonically-derived cell lines and the relative influence of CLAMP on splicing compared with transcription at target genes differs across in different cellular contexts.

### 3. CLAMP is highly enriched along gene bodies of sex-specifically spliced genes

What is the mechanism by which CLAMP regulates sex-specific splicing? If CLAMP directly regulates sex-specific splicing, we hypothesized that it would directly bind to DNA near the intron-exon boundaries of the genes where it regulates splicing. Therefore, we defined the binding pattern of CLAMP at the CLAMP-dependent female and male sex-specifically spliced genes in sexed embryos using CLAMP ChIP-seq data (#GSE133637). We found that 43.8% percent of all CLAMP-dependent sex-specifically spliced genes are bound by CLAMP across sexes and time points: 21.9% in 0-2 Hr female embryos, 8.2% in 0-2 Hr male embryos, 65.2% in 2-4 Hr female embryos and 59.4% in 2-4 Hr male embryos (**Supplementary Table S6**). The increase in percentage of genes bound by CLAMP in 2-4 Hr embryos compared with 0-2 Hr embryos is consistent with the known increased number and occupancy level of CLAMP binding sites at the later time point^29^. We also generated average profiles for CLAMP occupancy at genes where CLAMP regulates females (**red line**) and males (**blue lines**) sex-specific splicing in 0-2 Hr (pre-MZT) (**Fig 3A, B**) and 2-4 Hr (post-MZT) (**Fig 3C, D**) along both pre-and post-MZT time points in females (**Fig 3A, C**) and males (**Fig 3B, D**). We found that CLAMP occupies the gene bodies of many sex-specifically spliced genes that require CLAMP for their splicing. Overall, these data are consistent with a direct role for CLAMP in regulating splicing of sex-specifically spliced genes at up to 65% of its target genes.

**Fig 3.**
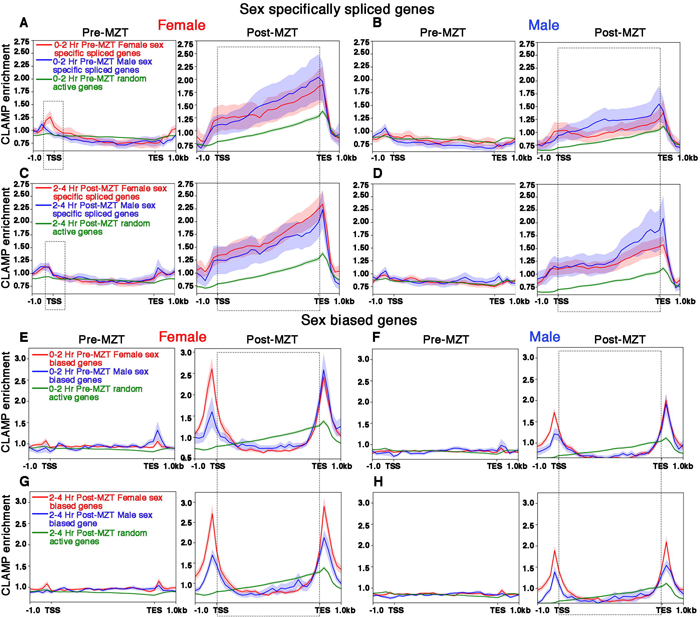
CLAMP binds along the gene body of female and male sex-specifically spliced genes at the post-MZT embryonic stage. **A-D** Average profiles for CLAMP binding at pre-MZT and post-MZT embryonic stages in females (**A, C**) and males (**B, D**) for genes **spliced** female-specifically (red line) and male-specifically (blue line) during the pre-MZT (**A, B**) and post-MZT (**C, D**) stages. **E-H** Average profiles for CLAMP binding to genes **expressed** in a sex-biased manner in females (red line) and males (blue line) during pre-MZT (**E, F**) and post-MZT (**G, H**) stage. Green lines in **A-H** represent CLAMP binding at a random set of active genes used as a control (see **Material and Methods** for details). Stippled regions in **A, C** (female, 0-2 Hr pre-MZT) denote chromatin around the TSS with more CLAMP binding in female sex-specifically spliced genes vs. male sex-specifically spliced genes. The dotted boxes in **A-H** highlight the gene body regions in CLAMP-dependent sex-specifically spliced genes and genes with CLAMP-dependent sex-biased expression.

Next, we compared the average CLAMP binding pattern at sex-specifically spliced genes (**Fig 3A-D**) to the CLAMP binding pattern at genes whose transcription but not splicing is both sex-biased and dependent on CLAMP (**Fig 3E-H**). In contrast to sex-specifically spliced genes where CLAMP occupancy is present over gene bodies, at genes that are expressed but not spliced in a CLAMP-dependent and sex-biased manner, CLAMP is enriched at the TSS and TES (area within the rectangular box demarcated by TSS and TES in **Fig 3A-H**). Furthermore, CLAMP binding is also modestly enriched at the TSS of female-biased expressed genes in females, consistent with enhanced CLAMP occupancy at the TSS of expressed genes ^40^. As a control, we used a random set of active genes that are not regulated by CLAMP (**green lines** in **Fig 3A-H**) and we observed lower occupancy than at CLAMP-dependent genes. Overall, we found preferential binding of CLAMP along the gene bodies of genes that have CLAMP-dependent splicing in both females and males in contrast to TSS and TES binding at genes where expression but not splicing requires CLAMP.

To determine whether the binding of CLAMP to gene bodies occurs close to splice junctions, we measured the distance between CLAMP peaks and the nearest splice junction (**Fig S6**). We found that CLAMP peaks are most frequently within 200-400bp of either the start or the end of a splice junction, especially in sex-specifically spliced genes. The resolution of these measurements is limited by sonication and therefore it is possible that binding occurs even closer to splice junctions. We also found that CLAMP binds to chromatin closer to splice junctions at CLAMP-dependent sex-specifically spliced genes compared to genes with CLAMP-dependent sex-biased transcription in 2-4 hr female embryo samples which have the most target genes and CLAMP binding events which improves statistical significance. The results were similar for all CLAMP peaks (**Fig S6C**) compared to peaks only present in introns (**Fig S6G**). Together these data support a direct role for CLAMP in modulating co-transcriptional RNA processing at a subset of its targets that we hypothesized is due to direct contact with target RNA transcripts and altering the recruitment of spliceosome components.

### 4. CLAMP binds to RNA on chromatin at a subset of sex-specifically spliced genes and interacts with the mature spliceosome complex sex-specifically

To test our hypothesis that CLAMP directly regulates co-transcriptional RNA splicing by contacting both the DNA and RNA of sex-specifically spliced genes and altering spliceosome recruitment, we asked whether CLAMP directly binds to RNA with fractionation iCLIP ^41, 42^ (**i**ndividual-nucleotide resolution **C**ross**L**inking and **I**mmuno**P**recipitation). We used cell lines for the iCLIP assay due to the large amount of input material required that could not be obtained from our low yield meiotic drive embryo system. Although CLAMP does not have a canonical **R**NA **r**ecognition **m**otif (**RRM**), it has a prion-like intrinsically disordered domain which mediates RNA interaction in many RNA binding proteins^43, 44^. Using single nucleotide resolution UV crosslinked immunoprecipitation (iCLIP) which defines direct protein-RNA interactions^41^, we determined that CLAMP binds directly to hundreds of RNAs and most targets are sex-specific with only 15% (124/816) of the target RNAs shared between male and female cell lines (**Fig 4A, Supplementary Table S7**). Also, we predicted unique CLAMP RNA binding motifs in male and female cells suggesting interaction with cofactors may change the ability of CLAMP to interact with RNA (**Fig S4A**).

**Fig 4.**
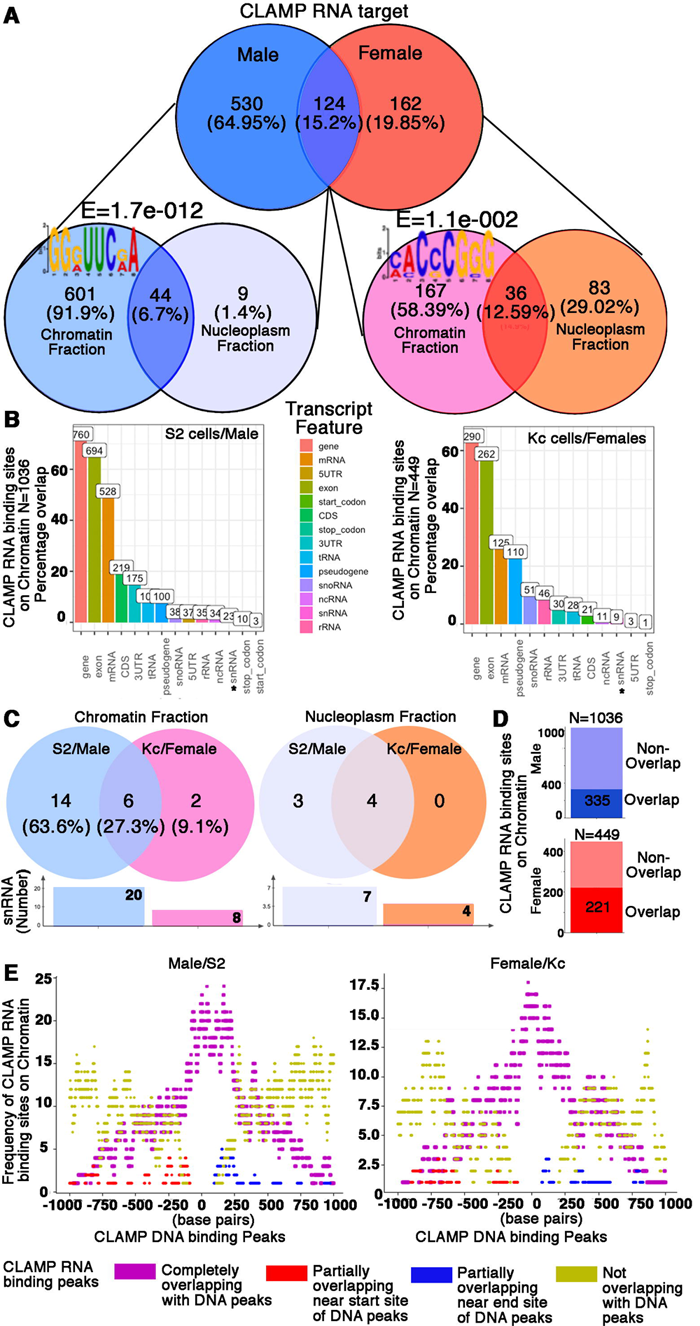
CLAMP binds to specific RNA transcripts on chromatin in males and females. **A.** Venn diagrams showing distribution of CLAMP RNA targets between male and female cell types and between chromatin and nucleoplasm fractions (**Four replicates** for each category performed except **3 replicates** for the Kc nucleoplasm fraction). **B.** Bar plots showing the percentage distribution of CLAMP RNA targets into different RNA categories in male (**S2**) and female (**Kc**) cell lines. The snRNAs, part of spliceosome complex is marked by asterisk on the x-axis. **C.** Venn diagrams showing distribution of CLAMP snRNA targets between male and female cell types in chromatin and nucleoplasm fractions. The corresponding bar plots denote the total number of snRNAs CLAMP binds to in respective fractions and cell types. **D.** Bar plots showing number of CLAMP RNA binding peaks overlapping with CLAMP DNA binding peaks in males (**blue bar**) and females (**red bar**). **E.** Frequency distribution of CLAMP RNA binding peaks (**iCLIP data, four replicates**) plotted over a region spanning CLAMP DNA binding peaks (**CUT&RUN data, 3 replicates for S2, 2 replicates for Kc**). Complete overlaps are denoted by magenta, non-overlaps in yellow, partial overlaps in red (near starting boundary of DNA peaks) and blue (near ending boundary of DNA peaks).

We identified CLAMP RNA binding targets separately in chromatin and nucleoplasmic cellular fractions to test whether CLAMP is binding to both DNA and RNA of target genes on chromatin. Also, identifying CLAMP RNA targets on chromatin allowed us to determine whether CLAMP is directly involved in co-transcriptional RNA processing at a subset of its targets. Interestingly, most CLAMP interaction with RNA occurs on chromatin (91.9%, 601/654 of male RNA targets; 58.4%, 167/286 of female RNA targets) (**Fig S4A**). Even though iCLIP was conducted in S2 and Kc embryonic cell lines, we still found 47/388 (**Column I, Supplementary Table S7**) target genes where CLAMP regulates sex-specific splicing in embryos and interacts with RNA by iCLIP in embryonic cell lines. Therefore, CLAMP sex-specifically and directly interacts with RNA targets on chromatin including the RNA encoded by a subset of the genes at which it regulates sex-specific splicing.

Next, we determined the overlap between genes whose splicing is regulated by CLAMP in larvae and cell lines and our cell line iCLIP data. In L3 larvae, 16 of the 124-female sex-specific and 140-male sex-specifically spliced CLAMP-dependent genes are direct CLAMP RNA targets identified from cell line iCLIP data (**Supplementary Table S3, Column I, Sheet2 and 3**), including key genes involved in sex-specific splicing: 1) the master sex-determination regulator and splicing factor *sxl;* 2) *hrp36*, an RBP that regulates alternative splicing; 3) *mrj,* which interacts with *hrp38* that regulates alternative splicing. The *hrp36* and *hrp38* genes are orthologous of each other and the well-studied human hnRNPA/B family of splicing factors^45^; 4) *bacc,* a splicing target we validated in embryos (**Supplementary Table S2**). Furthermore, we found that out of 615 splicing events (452 genes) that differed between Kc and S2 cells, 54 RNAs directly interact with CLAMP targets by iCLIP including key regulators of splicing (**Column I, Sheet 1, Supplementary Table S5 and Fig S7A**). When we compared the CLAMP-dependent splicing events in cell lines with direct CLAMP iCLIP targets, we identified 10 genes which are direct CLAMP RNA targets suggesting that a subset of the splicing events is regulated by direct contact between CLAMP and RNA (**Column I, Sheet 2&3, Supplementary Table S5 and Fig S7B**). Thus, many of the direct CLAMP splicing targets are key regulators of alternative splicing suggesting that CLAMP functions as an upstream master regulator of sex-specific splicing by directly regulating the splicing of a subset of key splicing factors which then regulate alternative splicing of additional indirect targets.

In addition to target RNAs that require CLAMP for their alternative splicing, CLAMP also directly interacts with spliceosomal RNAs sex-specifically. We found that CLAMP binds to more snRNAs in males (snRNA=20) including U1-U6 snRNAs (**Supplementary Table S7**) compared to females (snRNA=8) (**Fig 4B, C**). In the male chromatin fraction, CLAMP interacts with the catalytic step 2 spliceosome consisting of U2, U5, U6 snRNAs (FDR:1.7E-3). In contrast, the female chromatin fraction is enriched for transcripts that encode proteins that bind to the U1-U2 snRNAs (FDR:1.1E-2), suggesting a different type of regulation of splicing in males and females. Furthermore, we determined the overlap between CLAMP-RNA interaction sites in the chromatin fraction (iCLIP data) with CLAMP DNA binding peaks (CUT&RUN^46, 47^ **c**leavage **u**nder **t**argets and **r**elease **u**sing **n**uclease) from S2 (male) and Kc (female) cell lines that we generated (#GSE220053). We found that 32.3% (335/1036) of CLAMP RNA binding peaks in the S2 cell chromatin fraction and 49.2% (221/449) of CLAMP RNA binding peaks in the Kc cell chromatin fraction overlapped with CLAMP DNA binding peaks in the respective cell lines (**Fig 4D**). Next, we plotted the frequency of identifying a CLAMP RNA peak on chromatin over a region ±1 kb from the closest CLAMP DNA binding peak. We found that most overlapping CLAMP RNA peaks are within 250bp of the middle of the nearest CLAMP DNA peak in both male and female cells (**Fig 4E**), suggesting that CLAMP is linking RNA to DNA during co-transcriptional splicing at a subset of genes.

Integration of splicing analysis (**Figs 1, 2 & S2, 4/Tables S1, S3 and S5**), DNA-protein interaction data (ChIP-seq and CUT&RUN) (**Figs 3 & S6/Table S6**) and RNA-protein interaction data (iCLIP) (**Figs 4, S7/Table S7**), suggest that CLAMP interacts directly with a subset of its RNA targets to regulate co-transcriptional splicing, including key splicing regulators which likely amplify its function. Furthermore, CLAMP interacts with spliceosomal RNAs sex-specifically (**Fig 4B, C and Supplementary Table S7**). However, this integration does not fully explain how CLAMP regulates splicing in a sex-specific manner because CLAMP only binds to the RNA encoded by a subset of genes whose splicing is regulated by CLAMP. Therefore, we next asked how CLAMP modulates the function and localization of RBPs that that have known sex-specific roles including those that physically associate with CLAMP from proteomic studies^23^.

### 5. CLAMP interacts with RBP components of the spliceosome and influences their occupancy on chromatin

Because CLAMP differentially regulates splicing in male and female cells, we hypothesized that CLAMP regulates recruitment of RBP components of the splicing machinery to chromatin differentially in males and females. To test our hypothesis, we first examined how CLAMP regulates the occupancy of MLE, an RNA helicase that is a component of both the MSL complex, present only in males^48^, and the spliceosome, present in both sexes^49^. Previous studies showed that CLAMP physically associates with MLE^50–52^. Therefore, we hypothesized that CLAMP regulates the distribution of MLE between the spliceosome and the MSL complex to modulate sex-specific alternative splicing in males.

To test this idea, we measured MLE distribution across the genome using CUT&RUN^46, 47^. We performed CUT&RUN in both the presence and absence of maternal CLAMP, during both the pre-MZT and post-MZT embryonic stages in males (see methods). Our results show that MLE binds to chromatin both in males and females, with stronger binding in males (**Fig 5A**). Furthermore, the absence of CLAMP results in loss of MLE peaks in males but largely does not change MLE peaks in females (**Fig 5A, B**). This supports our hypothesis that CLAMP regulates MLE recruitment to chromatin in males.

**Fig 5.**
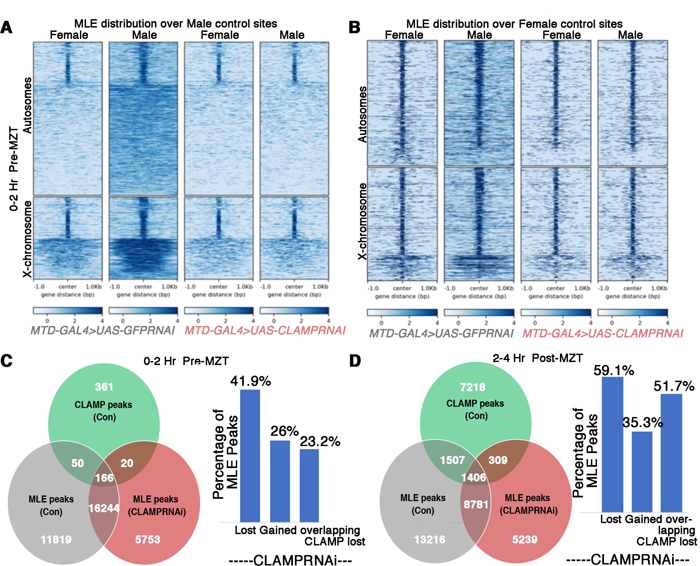
CLAMP regulates the distribution of MLE on chromatin in males. **A-B.** Heat maps showing the distribution of MLE at the male-specific (**A**) and female-specific (**B**) control MLE peaks on the X-chromosome and autosomes in male and female 0-2 Hr/pre-MZT embryos in the presence of maternal CLAMP (*MTD-GAL4>GFP RNAi*) and after the loss of maternal CLAMP (*MTD-GAL4>CLAMP RNAi)*. **C-D.** Venn diagrams and bar plots showing loss and gain of MLE peaks in the presence and absence of maternal CLAMP in male 0-2 Hr/pre-MZT (**B**) and 2-4 Hr/post-MZT (**C**) embryos. CLAMP peaks were identified only under control conditions (green circle), whereas MLE peaks were identified in the presence (grey circle) and absence (red circle) of maternal CLAMP protein depleted using the *MTD-GAL4>CLAMPRNAi* system.

We next compared the distribution and location of MLE peaks with that of CLAMP peaks previously identified in control embryos at the same time points^28^ and classified MLE peaks into two groups: 1) MLE peaks that overlap with CLAMP peaks (**Fig 5C,D and Fig S8**) and 2) MLE peaks that do not overlap with CLAMP peaks (**Fig 5C,D and Fig S8**). We found that MLE peaks that overlap with CLAMP peaks are largely at promoters in both developmental stages (**Fig S8**). In contrast, the MLE peaks that do not overlap with CLAMP peaks are primarily localized to introns (both X-chromosomal and autosomal peaks in both developmental stages (**Fig S8**)).

In the absence of CLAMP, there is a considerable loss and redistribution of both overlapping and non-overlapping MLE peaks in males. We found that overall 59.1% (14,723/24,910) of MLE peaks were lost at the 2-4 Hr/post-MZT stage in the absence of CLAMP. Moreover, ∼26% (5773/22183 pre-MZT) and ∼35% (5548/15735 post-MZT) of the MLE peaks observed in the absence of CLAMP were new peaks that were not present in control embryos (**Fig 5C-D**). After the loss of maternal CLAMP, ∼23% (50/216) of MLE peaks overlapping with CLAMP peaks are lost at the pre-MZT stage which increases to ∼51% (1,507/2,913) at the post-MZT stage (**Fig 5C-D**). Overall, our data suggest that MLE is redistributed in the absence of CLAMP. Therefore, CLAMP prevents aberrant recruitment of MLE in addition to the formation of aberrant splice isoforms (**Fig 2C**). Furthermore, we hypothesize that MLE which is at the new peaks in the absence of CLAMP is part of the spliceosome complex and not MSL complex because MSL complex is not present on chromatin in the absence of CLAMP^40^.

To provide insight into the differences between MLE peaks which overlap with CLAMP and those which do not, we identified sequence motifs which are enriched within each class of peaks using MEME within the time2splice pipeline. The known CLAMP motif ^40^, a stretch of (GA)n repeats, is enriched at regions that are bound by both MLE and CLAMP independent of stage and chromosome type as expected. In contrast, MLE peaks which do not overlap with CLAMP have motifs with stretches of GTs, GCTs, and GTAs but not (GA)n repeats (**Fig S8**). In the absence of CLAMP, the remaining MLE peaks (red circle) were most enriched for (GT)n motifs (**Fig S8C, D**) which have known roles in splicing through encoding secondary RNA structures^53–55^. Therefore, CLAMP prevents MLE from redistributing to sequence motifs that are known regulators of splicing.

We also found that CLAMP changes the distribution of MLE relative to genes, increasing the frequency at which MLE is present at promoters instead of introns. MLE peaks that overlap with CLAMP peaks (intersection of green circle with red and grey circles, **Fig S8**) on the X-chromosome (**Fig S8A, C**) or autosomes (**Fig S8B, D)** are enriched at promoters (**Fig S8A, C** (X-chromosome), **Fig S8B, D** (Autosomes)). In contrast, **new** unique MLE peaks not overlapping with CLAMP (grey area in Venn diagrams, **Fig S8**) and those that are gained after *clamp* RNAi (red area in Venn diagrams, **Fig S8**) are enriched at introns (**Fig S8A, C** (X-chromosome), **Fig S8B, D** (Autosomes)). These results suggest that CLAMP sequesters MLE at (GA)n rich sequences within promoters that prevents it from binding to GT motifs within introns which regulate splicing^53–55^. Thus, in the absence of CLAMP, MLE is redistributed and aberrantly binds to intronic sequences with known motifs that regulate splicing independent of whether present on the X-chromosome or autosomes.

To determine how MLE redistribution could alter sex-specific splicing, we plotted the distribution of MLE binding on CLAMP-dependent female and male-specifically spliced genes in the presence and absence of CLAMP (**Fig S9A, B**). Pre-MZT, MLE binds near the TSS of male-specifically spliced genes independent of maternal CLAMP (**Fig S9A**). At the post-MZT stage, loss of maternal CLAMP in male embryos causes MLE to change its binding distribution along the gene body (rectangle with dotted lines: **Fig S9B**) of CLAMP-dependent male-specifically spliced genes (blue line) relative to CLAMP-dependent female-specifically spliced genes (red line). These profiles are consistent with a model in which CLAMP regulates MLE distribution at male-specifically spliced genes to alter male sex-specific splicing. In males without CLAMP, increased MLE binding at female sex-specifically spliced genes (red line, enclosed within rectangle with dotted lines: **Fig S9B**) may result in aberrant differential splicing of these genes. Thus, our data suggests that CLAMP inhibits mis-localization of MLE to female sex-specifically spliced genes in males.

Next, we asked whether CLAMP associates with spliceosome complex protein components other than MLE, which is a component shared with the MSL complex. We have shown that CLAMP directly binds to snRNAs (**Fig 4C**) and previously reported that CLAMP physically associates with several spliceosome complex components based on mass spectrometry analysis^23^. To validate these associations, we performed co-immunoprecipitation (coIP) experiments to measure association between CLAMP and two hnRNP spliceosome components with known functions in sex-specific splicing, the conserved hrb27C and Squid proteins ^45, 49, 56^. We found that in both S2 (male) and Kc (female) cells, CLAMP interacts with hrb27C. In contrast, CLAMP only associates with Squid in female Kc cells and not in male S2 cells (**Fig S10A, B**), consistent with mass spectrometry data. In contrast to MLE and CLAMP which are enriched on the male X-chromosome, we found that Squid occupancy on polytene chromosomes is decreased on the male X-chromosome compared with the female X chromosome (**Fig S10C-E**). Therefore, it is possible that there is a competition between CLAMP recruitment of MSL complex to the male X-chromosome and CLAMP recruitment of the spliceosome complex containing Squid that contributes to sex-specific splicing. Overall, CLAMP differentially associates with RBP spliceosome components in males and females, providing a potential mechanism by which CLAMP can regulate sex-specific splicing.

### 6. CLAMP regulates the chromatin accessibility and splicing of the sxl gene, directly interacts with sxl RNA, and alters splicing of other sex determination pathway component genes

Next, we asked how CLAMP physically and functionally interacts with known key regulators of sex-specific splicing. In *Drosophila*, sex-specific alternative splicing is regulated by the sex-determination pathway. Sex-lethal (Sxl) is the master regulator of sex determination^57^ and drives subsequent sex-specific splicing in females^58^ of downstream effector genes giving rise to female specific effector proteins (Fig 6A) that regulate female-specific splicing. Functional Sxl protein is only produced in females^57, 59^ because exon three in the *sxl* transcript contains a premature stop codon which is spliced out in females but retained in males ^60^. Absence of functional Sxl protein in males results in formation of male-specific effector proteins that regulate male-specific splicing (Fig 6A). Therefore, we asked whether CLAMP regulates alternative splicing of the *sxl* transcript.

**Fig 6.**
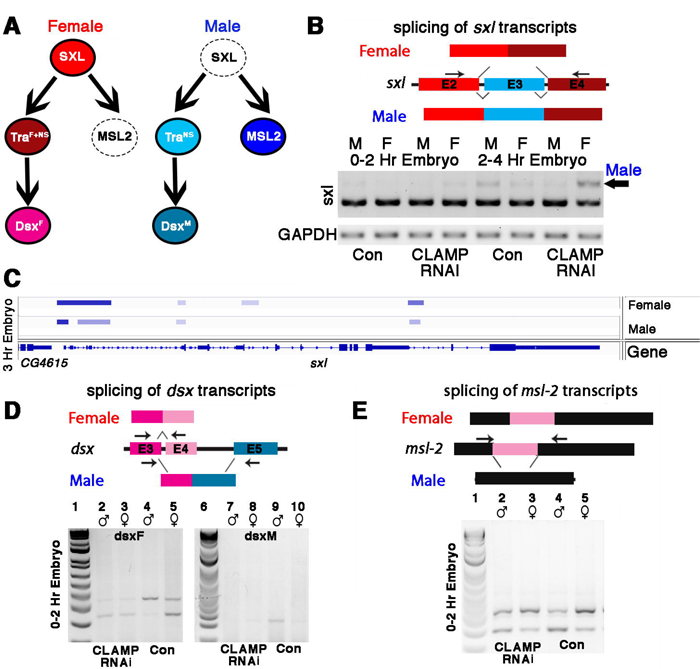
Alternative splicing of components of the sex determination pathway is regulated by maternal CLAMP in females. **A.** The sex determination pathway in *Drosophila* is regulated by master regulator Sex Lethal (SXL). **B.** RT-PCR electrophoresis gel images (inverted colors) showing splicing of *sxl* transcripts in 0-2 and 2-4 Hr sexed embryos in the presence and absence of maternal CLAMP with a representative schematic of the splicing event at the top of the gel image. The arrow indicates the male-specific *sxl* transcript. **(Number of replicates=2)** **C.** IGV browser image showing CLAMP ChIP-seq peaks (rectangular boxes in light blue) at the genomic locus for the *sxl* gene in male and female 3Hr embryos. For each sample, the narrow peak file is shown which is generated after peak calling. **D-E.** RT-PCR electrophoresis gel images from 0-2 Hr embryonic RNA samples (lane 2-5 & 7-10) showing splicing of *dsx* (**D**) and *msl2* (**E**) transcripts in females (lanes 3,5,8,10) and males (lanes 2,4,7,9). Embryos were laid by *MTD-GAL4>GFP RNAi* control (lanes 4,5,9,10) and *MTD-GAL4>CLAMP RNAi* (lanes 2,3,7,8) females. The schematic above each gel image shows the female and male splice variants of the *dsx* (**D**) and *msl2* (**E**) transcripts.

To test whether CLAMP regulates *sxl* alternative splicing, we designed an RT-PCR assay to distinguish between the female-specific (excluding exon 3) and male-specific (including exon 3) versions of the *sxl* transcript (**Fig 6B**). To determine whether maternal CLAMP regulates splicing of the *sxl* transcript, we performed RT-PCR analysis of *sxl* splicing. In contrast to the much later larval stage (**Fig S11A**), in embryos, the male and female isoforms of Sxl have not become fully specified, consistent with the known autoregulation of *sxl* that occurs in embryos^57, 59, 61, 62^. Despite the lack of complete specification of male and female *sxl* transcripts, our data show that maternal CLAMP promotes the sex-specific splicing of *sxl* transcripts in 0-2 and 2-4 Hr embryos because the male-specific transcript is not expressed in maternal CLAMP-depleted male embryos but is expressed in CLAMP-depleted female embryos (**Fig 6B**). Consistent with the ChIP-seq binding pattern of CLAMP at the *sxl* locus on chromatin (**Fig 3, 6C**) which show enhanced binding in 2-4 Hr embryos compared with pre-MZT embryos, there is an enhanced function for CLAMP in splicing at 2-4 hours compared with 0-2 hours.

Next, we assayed the function of zygotic CLAMP in *sxl* splicing in three previously described fly lines: 1) our recessive *clamp* null mutant *clamp*^2^ line ^27^; 2) the heterozygous mutant *clamp*^2^*/CyO-GFP* line; 3) our rescue line which is homozygous for the *clamp*^2^ allele and contains a rescue construct which is an insertion of the wild type *CLAMP* gene at an ectopic genomic location. We measured CLAMP-dependent changes in alternative splicing of *sxl* and found that in homozygous *clamp*^2^ female animals, there is a small but detectable amount of the longer male-specific *sxl* transcript (**Fig S11A**, lane c). This mis-regulation of *sxl* splicing is rescued by our *CLAMP*-containing rescue construct (**Fig S11A**, lane d). Furthermore, our iCLIP data show that CLAMP directly binds to *sxl* transcripts in female but not male cells (**Supplementary Table S7**) and our L3 RNA-seq data demonstrate that CLAMP regulates *sxl* splicing in females (**Supplementary Table S3**) and not *sxl* transcription (**Supplementary Table S4**). These data suggest that one way in which CLAMP functions in sex-specific splicing in females is upstream of Sxl by binding to both DNA and RNA at the *sxl* locus to regulate splicing.

To test whether defects in *sxl* splicing altered Sxl protein levels, we performed western blots to quantify Sxl protein in wild type females and males and *clamp*^2^ null females (**Fig S11B**). We observed a reduction in Sxl protein levels in females in the *clamp*^2^ null background when compared with controls. Also, homozygous *clamp*^2^ mutant males die before the late third instar larval stage, and therefore it was not possible to measure the splicing of transcripts in male *clamp*^2^ mutant larvae.

When comparing our RT-PCR assay measuring *sxl* splicing (**Fig S11A**) with western blotting analysis measuring Sxl protein levels (**Fig S11B**), we observed a more dramatic reduction in Sxl protein levels compared to changes in splicing. We have also found that CLAMP binds to the 5’UTR region of the *sxl* transcript in females (**Fig S11C**) and regulation of translation by 5’UTR binding is a common mechanism for regulating protein stability^63, 64^. Therefore, we speculate that CLAMP binding to the 5’UTR of *sxl* transcripts in females (**Supplementary Table S7**) may function in translational regulation of the Sxl protein. Furthermore, CLAMP interacts sex-specifically with the translation factor FMRP in the male cytoplasm (**Fig S11D**), indicating CLAMP might also have a distinct differential influence on translation in male and females depending on interacting translation regulatory proteins. Together, these data suggest that it is possible that mis-regulation of translation amplifies the CLAMP-dependent mis-regulation of splicing to generate a larger decrease in Sxl protein levels in the absence of CLAMP. Future experiments are required to test this hypothesis and decipher underlying mechanisms. Independent of a potential effect on translation mediated by sex-specific FMRP interaction, we determined that CLAMP promotes female-specific splicing of the *sxl* transcript as one mechanism to ensure that normal Sxl protein levels are produced in females.

Next, we wanted to determine the mechanism by which CLAMP regulates splicing of *sxl*. Recent reports provide strong evidence that increased chromatin accessibility contributes substantially to the retention of introns during AS^65^. In addition, splicing-associated chromatin signatures have recently been identified^66^. CLAMP regulates chromatin accessibility^29^ and both CLAMP ChIP-seq data from sexed embryos (**Fig 6C**) and CUT&RUN data from cell lines (**Fig S11E**) shows that CLAMP binds differentially to the *sxl* gene in females compared to males. Therefore, we measured chromatin accessibility at the *sxl* gene locus in S2 (male) and Kc (female) cells in the presence and absence of CLAMP by mining our previously generated Micrococcal Nuclease sequencing data^21^. Regions of the genome that are accessible have a positive MACC score (shown in blue), and regions of the genome that are inaccessible have a negative score (shown in red) (range is between -0.33(red) and +1.33 (blue)). We found that after the loss of CLAMP in female Kc cells, chromatin accessibility at exon three of *sxl* increases significantly (**Fig S11F**). In females in which CLAMP has been deleted but not control females, *sxl* exon three shows a strong and statistically significant MACC peak^21^ indicative of open chromatin (**Fig S11F** boxed rectangular inset). Therefore, CLAMP normally promotes a closed chromatin environment at exon three in females but not males.

Our MNAse-seq data combined with our splicing results and recent literature on the link between chromatin and splicing suggest that increased chromatin accessibility in males compared to females may cause the retention of exon three in the male *sxl* transcript. Consistent with our results in males, open chromatin marks such as H3K4me1& H3K4me2 are enriched just upstream of the start site of retained exons^66^. In contrast, histone marks associated with condensed chromatin such as H4K20me1&2, H3K9me3, and H3K27me3 are highly enriched at excluded exons^66^, consistent with our results in females.

To further define how CLAMP and Sxl function dependently and independently of each other to regulate sex-specific splicing, we asked whether CLAMP directly binds to RNA targets that have Sxl binding motifs using our iCLIP data. Overall, we found that ∼20% of CLAMP RNA targets have Sxl motifs suggesting that CLAMP and Sxl have shared and independent targets (**Fig S12A**). Next, we compared CLAMP RNA targets (nuclear iCLIP) from female Kc cells with the only available data identifying RNAs that interact with Sxl which was generated from adult females (#**GSE98187**). Even though the data sets are not from the same type of biological sample, we still determined that 81/286 (28.32%) of CLAMP bound RNAs in Kc cells overlap with Sxl targets in adult females (**Fig S12B, C and Supplementary Table S8**). Overlapping targets include *sxl* transcripts and snRNAU5, a component of the U5snRNP complex involved in splicing^67^. Interestingly, 26.5% (173/654) of CLAMP iCLIP RNA targets from S2 cells (male) overlapped with Sxl female RNA targets, indicating that CLAMP may bind to RNAs in male cells that are Sxl targets in females (**Fig S12B, C and Supplementary Table S8**).

Also, 88/388 (22.68%) of the CLAMP-dependent sex-specific spliced genes are Sxl targets (**Supplementary Table S9**). We also determined which of the CLAMP-dependent sex-specifically spliced genes products directly interact with CLAMP or Sxl or both factors (**Supplementary Table S10, Fig S12D**). These results further support our hypothesis that CLAMP functions through multiple mechanisms to regulate sex-specific splicing: 1) CLAMP regulates a subset of targets through direct binding to DNA and RNA of target genes including the Sxl gene itself and other key regulators of alternative splicing; 2) CLAMP regulates other targets indirectly through regulating Sxl splicing and protein levels.

To further understand the direct and indirect roles of maternal and zygotic CLAMP in sex-specific splicing, we examined the splicing of other components of the sex determination pathway downstream of Sxl (**Fig 6D, E and S11G-H**). In embryos which lack maternal CLAMP (**Fig 6D**, lane 2-3), the *dsx* female-specific transcript is aberrantly produced in males (**Fig 5D**, lanes 2-5). In contrast, the male-specific *dsx* transcript is not expressed in male embryos which lack CLAMP, similar to wild type female embryos (**Fig 6D**, lane7-10). We also observed male-specific *dsx* transcripts in female *clamp*^2^ mutant larvae (**Fig S11G**, lane c). Therefore, *dsx* splicing is regulated by maternal and zygotic CLAMP and CLAMP binds directly to the *dsx* gene locus (**Fig S12E**) but not to *dsx* RNA (**Supplementary Table 7**). These data suggest that CLAMP may regulate *dsx* splicing via both Sxl-dependent and Sxl-independent mechanisms.

In addition, we found that maternal and zygotic CLAMP regulates splicing of the male-specific lethal-2 (*msl-2*) transcript, which is male-specific because Sxl regulates its splicing, transcript stability, mRNA export, and translation in females^68^ (**Fig 6E, Fig S11H, lane c**). To determine whether splicing defects also cause dysregulation of MSL-2 protein expression and localization to chromatin, we performed polytene immunostaining from female *clamp*^2^ mutant salivary glands. In the absence of CLAMP, ectopic MSL2 protein (in red) is present at several locations on female chromatin in contrast to controls (*clamp*^2^*/CyO-GFP* heterozygous females) where MSL-2 protein is not present on chromatin, consistent with lack of MSL complex formation in wild type females (**Fig S11I**). Similar to *dsx*, the *msl-2* gene is also bound by CLAMP (**Fig S12F**) and regulated by Sxl and therefore could be regulated through both direct and indirect mechanisms. In addition, CLAMP binds to the DNA and RNA that encodes another sex-specific splicing regulator and Sxl target gene *fru* (*fruitless*) (**Supplementary Table S7**). Together, these data reveal that CLAMP regulates the splicing and protein expression of multiple components of the sex determination pathway via Sxl-dependent and Sxl-independent mechanisms.

To determine which factors may function with CLAMP to regulate sex-specific splicing in addition to Sxl, we analyzed our iCLIP CLAMP RNA binding data (**Supplementary Table 7**) for motifs of other RNA binding proteins involved in splicing^49^ (**Fig S13**). We found that 70-80% of CLAMP bound RNA sequences contain motifs for Tra which functions downstream of Sxl (Male Chromatin Fraction, **N=645**; Male Nucleoplasmic Fraction, **N=53**; Female Chromatin Fraction, **N=203**; Female; Nucleoplasmic Fraction, **N=119**). Furthermore, 30-50% of CLAMP bound RNA sequences contain motifs for Lark, a splicing factor homologous to human RBM4^69^, and *Drosophila* homologues of the hnRNPA/B family of splicing factors, Hrb98DE/Hrb87F/Hrb27C^49^ (**Fig S13**). We had previously identified an association of CLAMP with Hrb27C by MALDI-MS^23^ which we now validated by co-immunoprecipitation (**Fig S10A**). Together, our generation and integration of functional sex-specific splicing analysis with RNA-TF interactions, DNA-TF interactions, chromatin accessibility, and protein-protein interaction data reveal new mechanisms by which TFs function with RBPs to regulate co-transcriptional splicing that promotes a key developmental decision very early in development.

## Discussion

### A maternal factor links transcription to splicing during the earliest stages of sexual differentiation

Alternative splicing (AS) is a highly conserved mechanism that generates transcript and protein diversity^70–72^. Several studies have reported highly dynamic RNA bound proteomes (RBPs) during the Maternal Zygotic Transition (MZT) across diverse phyla, with widespread alternative splicing events occurring during early embryonic development^31, 73–76^. Furthermore, diverse isoforms are present in the pool of maternal and zygotic transcripts during early development ^30, 31^. However, the mechanisms that integrate the function of TFs and RBPs to regulate transcript diversity in a context-specific manner during the earliest hours of development remain elusive.

Maternally-deposited pioneer transcription factors drive zygotic genome activation, but their role in generating transcription diversity in the early embryo was unknown. Here, we define sex-specific alternatively spliced isoforms in pre-and post-MZT *Drosophila melanogaster* female and male embryos genome-wide for the first time. We show that sex-specific transcript diversity occurs much earlier in development than previously thought by generating the earliest data that define sex-specific transcript diversity across species. Furthermore, we identify how a maternally-deposited pioneer TF, CLAMP, regulates sex-specific transcript diversity in early embryos. Prior work on sex-specific transcript diversity^37, 56, 73, 76–82^ either examined sex-biased differences in gene expression only or sex-specific transcript diversity much later in development in adult gonads or brain. To overcome the challenge of sexing early embryos before zygotic genome activation, we used a meiotic drive system that generates sperm with either only X or only Y chromosomes^16^ and measured both transcription and sex-specific transcript diversity generated by alternative splicing.

We show even following the initial few hours of its existence, there is a clear difference between a male and female *Drosophila* embryo’s transcript variation that was not previously identified (**Figs 1, 2**). Because the transcript variants in both males and females encode genes that are involved in developmental processes, sex-specific developmental distinctions may occur earlier than previously thought. We demonstrate that a fundamental developmental trajectory differs between males and females from the initial hours of their existence long before gonad formation. Such early sex-specific transcript diversity may provide insight into how developmental disorders that originate before gonad formation can exhibit variable penetrance between sexes.

Different splice variants are produced at different frequencies over time and between sexes. To date, we lacked pipelines to characterize how these isoforms change over time. Therefore, we developed time2splice, which identifies mechanisms to regulate temporal and sex-specific alternative splicing by combining RNA-seq and protein-DNA interaction data from CUT&RUN and ChIP-seq experiments. Time2splice has three parts: 1) temporal splicing analysis based on the SUPPA algorithm; 2) temporal protein-DNA analysis, and 3) temporal multi-omics integration. The pipeline and analysis steps can be accessed at: https://github.com/ashleymaeconard/time2splice.

We defined groups of genes in both males and females that undergo alternative splicing events which are regulated by maternally-deposited and zygotically-expressed CLAMP. Thus, the maternal environment both regulates transcription initiation and shapes RNA processing events that are maintained later during development. The key question is: How does CLAMP, a ubiquitously expressed pioneer TF, regulate sex-specific splicing? We identified several mechanisms by which CLAMP regulates sex-specific splicing.

### CLAMP regulates sex-specific splicing via multiple mechanisms that include context-specific interaction with target RNAs and RBPs

CLAMP binds directly to intronic regions of approximately half of the sex-specifically spliced genes that it regulates in both males and females suggesting a direct role in regulating their co-transcriptional splicing by altering the recruitment of spliceosomal components or chromatin accessibility. Our data supports a model in which direct CLAMP binding to DNA and RNA regulates the splicing of a subset of its target genes (**Fig 3 & 4D, E and Supplementary Table S6&S7**). Many of these direct target genes are key regulators of alternative splicing, further enhancing the effect of CLAMP on splicing (**Supplementary Table S2, S7**). Because not all CLAMP-interacting RNAs on chromatin are differentially spliced, we hypothesize that CLAMP has other regulatory co-transcriptional functions such as potentially regulating transcript stability or nuclear export which require future investigation.

Furthermore, CLAMP regulates chromatin as a pioneer TF^21, 29^ and recent literature links chromatin and splicing^65, 66^. For example, closed chromatin marks have recently been linked to exon exclusion and open chromatin has been linked to exon inclusion^65, 66^. Our results also indicate that CLAMP associates with the functional spliceosome complex in males but not in females (**Supplementary Table S7 and Fig 4C**). Proteomic analysis^23^ and coIPs (**Fig S10 A, B**) show that CLAMP is associated with spliceosome complex components like Squid, a known to regulate sex-specific splicing^56^, specifically in females and with MLE, a component of both the spliceosome and MSL complex^48^ only in males. These data support a model in which differential association between CLAMP and RBP spliceosome complex components in males and females regulates sex-specific splicing. Thus, we hypothesize that CLAMP may recruit RBP spliceosome complex components to regulate splicing by altering the chromatin environment or/and directly binding to target RNA transcripts (**Fig 7A**).

**Fig 7.**
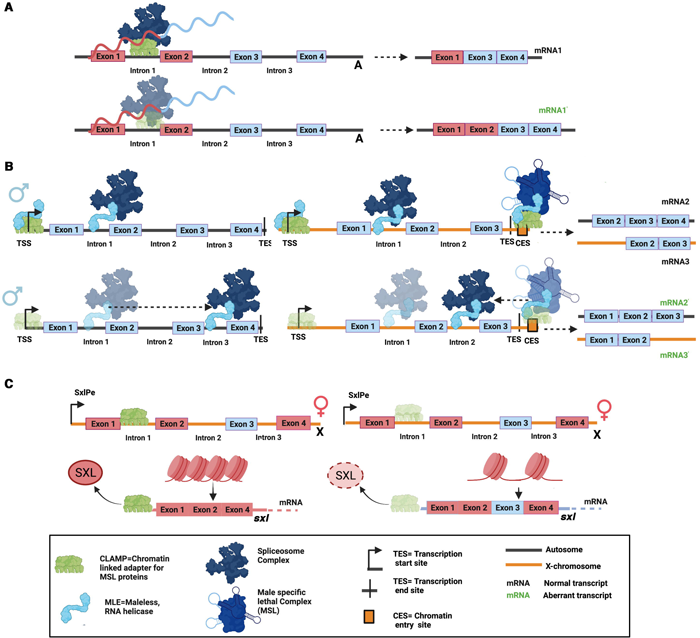
Mechanisms by which CLAMP regulates sex-specific splicing in females and males. **A.** CLAMP regulates splicing in both males and females via directly binding to intronic DNA sequences of CLAMP-dependent sex-specifically spliced genes and sex-specific interaction with a subset of sex-specifically spliced RNAs and sex-specific interaction with spliceosomal RNAs. **B.** CLAMP may regulate the distribution of MLE between the spliceosome and the male X-chromosome specific MSL complex in males. CLAMP increases the occupancy of MLE at promoters and CES. In the absence of CLAMP, MLE is lost from many of its binding sites, including CES and promoters, and is gained at ectopic intronic sequences which contain motifs that regulate splicing which correlates with aberrant sex-specific splicing in males. **C.** In females, CLAMP binds near the SxlPe promoter and regulates chromatin accessibility at exon three (blue square) of the *sxl* gene and binds to the *sxl* mRNA. In this way, CLAMP promotes the excision of exon3 such that functional Sxl protein is formed, which drives female-specific splicing events. The absence of CLAMP in females thus results in the aberrant production of non-functional male-specific *sxl* transcripts which retain exon3, reducing levels of functional Sxl protein. CLAMP also binds to the 5’UTR of the *sxl* RNA which may regulate its export or translation. CLAMP and Sxl have shared and distinct RNA targets suggesting that they function by both dependent and independent mechanisms. The three mechanisms proposed in parts **A**, **B**, and **C** are not mutually exclusive and are likely to occur simultaneously.

Our results also show that CLAMP inhibits aberrant splicing events in males, especially at the post-MZT stage (**Fig 2C**) and the distribution of MLE, an RNA helicase component of the spliceosome on chromatin is CLAMP-dependent (**Fig 4A**). = In the absence of CLAMP, the fraction of promoter bound MLE is reduced and MLE re-localizes from its normal intronic binding sites to new intronic regions that contain GT sequence motifs (**Fig S8**) known to regulate splicing^53–55^. Therefore, we hypothesize that CLAMP regulates the localization of MLE to suppress aberrant female-specific splicing events in males. Because MLE is part of the MSL complex only in males and the spliceosome complex in both sexes, we hypothesize that CLAMP influences the relative distribution of MLE between two different ribonucleoprotein complexes: 1) the MSL complex; and 2) the spliceosome, co-regulating sex-specific splicing and male X-chromosome dosage compensation (**Fig 7B**). Without CLAMP, the MSL complex does not localize to the X-chromosome and becomes destabilized ^40^; thus, MLE is no longer part of the MSL complex and is available to redistribute to new spliceosome binding sites. Therefore, we provide evidence to support a model in which CLAMP sex-specifically inhibits aberrant binding of MLE to motifs that regulate splicing which alters sex-specific transcript diversity.

To provide mechanistic insight into how a pioneer transcription factor like CLAMP regulates sex-specific splicing in females, we investigated the role of CLAMP in regulating the *sxl* gene locus, the master regulator of sex-determination pathway^57^. CLAMP binds near the early promoter of the *sxl* gene (SxlPe) and regulates the chromatin environment at exon three of *sxl* which is normally spliced out in females (**Fig 7C**). Consistent with recent literature linking chromatin accessibility to alternative splicing^65, 66^, we hypothesize that closed chromatin at exon three induces exclusion of this exon from female *sxl* transcripts. In the absence of CLAMP in females, the chromatin becomes more open, and the *sxl* transcript is not bound by CLAMP. Therefore, exon three is included in a subset of *sxl* transcripts in females which reduces the levels of functional Sxl protein due to the incorporation of a stop codon, thus dysregulating downstream splicing events (**Fig 7C**). Because CLAMP binding sites are present near the promoter region of the *sxl* gene, we hypothesize that CLAMP regulates chromatin at exon three from a distance, consistent with our recent findings suggesting that CLAMP can mediate long-range chromatin interactions ^83, 84^ and act on chromatin accessibility at a distance ^21^.

Furthermore, CLAMP binds to the 5’UTR of the *sxl* transcript (**Fig S11H**) specifically in females, which we hypothesize is important for regulating *sxl* translation because we observe a stronger reduction in Sxl protein levels in females in absence of CLAMP compared with the effects on female-specific splicing (**Fig S11A, B**). Both the decreased Sxl protein levels in female *clamp*^2^ mutants and mis-expression of female and male-specific *dsx* transcripts suggest that CLAMP may regulate sexual differentiation because sex-specific Dsx protein isoforms are known determinants of sexual dimorphism across species^57^. Also, CLAMP directly binds to the DNA of *sxl, dsx,* and *msl-2* target genes, suggesting a direct role in regulating these loci. Furthermore, CLAMP binds to the DNA and RNA of *sxl* and *fru*. Fruitless (*fru*) encodes a BTB zinc finger transcription factor that contributes to sexual differentiation of neural circuits ^85, 86^ and many CLAMP-dependent sex-specifically spliced genes regulate neural development (**Fig 2E**).

Because CLAMP and Sxl have both overlapping and distinct targets (**Supplementary Tables S8, 9, 10 and Fig S13A-D**), we hypothesize that CLAMP regulates sex-specific splicing both via the Sxl-mediated sex-determination pathway as well as independent from Sxl. In support of our hypothesis, CLAMP RNA binding sequences share motifs with other RNA binding proteins involved in splicing (**Fig S13E**), some of which are sex-specific protein interaction partners of CLAMP^23^. Therefore, CLAMP may regulate splicing through at least two possible mechanisms that are not mutually exclusive: 1) CLAMP directly regulates the splicing of a subset of sex-specifically spliced genes by linking RNA to chromatin and altering the recruitment of the spliceosome; 2) CLAMP regulates the sex-specific splicing of transcripts indirectly by functioning upstream of Sxl which is a known regulator of splicing of downstream genes such as *msl-2* and *dsx*.

Overall, we hypothesize that both different composition of the spliceosome and its differential recruitment to chromatin drive sex-specific changes in splicing. We identify CLAMP as a maternal factor that regulates sex-specific alternative splicing through its sex-biased association with the DNA and RNA of target genes, sex-biased recruitment of spliceosome components, and its ability to influence the sex determination pathway. Identifying the factors that regulate this sex-biased association of CLAMP with spliceosome complex components will be a key future direction.

Here, we show for the first time that a maternal factor controls sex-specific splicing during early embryonic development, highlighting how the maternal environment influences transcript diversity in the zygote from activation of the zygotic genome to the processing of zygotic RNA products. Consistent with recent literature linking chromatin accessibility and splicing, our results suggest that CLAMP could be one example of a more general splicing regulatory mechanism controlled by the interaction between pioneer TFs that alter chromatin accessibility and components of the RNA processing machinery which generates spatial-temporal transcript diversity. Consistent with this hypothesis, many transcription factors have recently been shown to interact directly with RNA^87–89^. While we analyzed sex-specific transcriptome diversity in this study and linked it to the sex-specific dosage compensation process, similar mechanisms could drive cell-type specific variation. For example, cell fate-determining transcription factors could regulate the chromatin occupancy of splicing complex components to promote the formation of cell-type-specific isoforms. We also present time2splice, a new pipeline to uncover mechanisms which drive such spatial-temporal transcript diversity by integrating splicing and chromatin occupancy data.

## Materials and Methods

### Fly strains and husbandry

*Drosophila melanogaster* fly stocks were maintained at 24°C on standard corn flour sucrose media. Fly strains used: *MTD-GAL4* (Bloomington, #31777), *UAS-CLAMPRNAi[val22]* (Bloomington, #57008), Meiotic drive fly stocks +; SD72/CyO and 19-3, yw, Rsp[s]-B[s]/Dp(2:y)CB25-4, y+, Rsp[s]B[s]; SPSD/CyO (Bloomington, #64332) (both gifts from Cynthia Staber). These were crossed to obtained male and female embryo of desired genotypes according to Rieder et al 2017.

### Cell culture

Kc and S2 cells were maintained at 25°C in Schneider’s media supplemented with 10% Fetal Bovine Serum and 1.4X Antibiotic-Antimycotic (Thermofisher Scientific, USA). Cells were passaged every 3 days to maintain an appropriate cell density.

### Sample collection and Western blotting

Salivary glands from third instar larvae were dissected in cold PBS and samples frozen in liquid nitrogen. Total protein from the samples was extracted by homogenizing tissue in the lysis buffer (50mM Tris-HCl pH 8.0, 150mM NaCl, 1% SDS, 0.5X protease inhibitor) using a small pestle. After a five-minute incubation at room temperature, cleared the samples by centrifuging at room temperature for 10 minutes at 14,000xg. To blot for CLAMP and Actin, 5 micrograms of total protein was run on a Novex 10% Tris-Glycine precast gel (Life technologies). To measure Sex-lethal protein levels, 20 micrograms of total protein was run on a Novex 12% Tris-Glycine precast gel (Life technologies). Protein was transferred to PVDF membranes using the iBlot transfer system (ThermoFisher Scientific) and probed the membranes for CLAMP (1:1000, SDIX), Actin (1:400,000, Millipore), and SXL (1:500, a gift from Fatima Gebauer) antibodies using the Western Breeze kit following the manufacturer’s protocol (ThermoFisher Scientific). We quantified the relative expression of protein for SXL using the gel analysis tool in ImageJ software following the website’s guidelines ^90^. For each genotype, we first internally normalized the amount of SXL protein to Actin. Next, we determined the protein’s relative expression by comparing the Actin normalized quantities to y[1], w[1118] female samples.

### Polytene chromosome squashes and immunostaining

Polytene chromosome squashes were prepared as previously described in Reider et al. 2017. We stained polytene chromosomes with rabbit anti-CLAMP (1:1000, SDIX), mouse anti-Squid (1:50, 1B11, DSHB), rat anti-MSL2 (1:500, gift from Peter Becker) antibodies. For detection, we used all Alexa Fluor secondary antibodies against rabbit and mouse at a concentration of 1:1000 and visualized slides at 40X on a Zeiss Axioimager M1 Epifluorescence upright microscope with the AxioVision version 4.8.2 software.

### Splicing assays for male and female-specific transcripts

To test for the male and female splice forms of *sex-lethal, transformer, doublesex*, and *msl2*, total RNA was extracted from ten third instar larvae from each genotype. We reverse-transcribed two micrograms of total RNA using the SuperScript VILO cDNA Synthesis Kit (Life Technologies) following the manufacturer’s protocol. We amplified target sequences by PCR using primers designed to span Alternatively spliced junctions. Alternative splicing primer sequences for sxl FP- TGCAACTCACCTCATCATCC, sxl RP- GATGGCAGAGAATGGGACAT, for tra FP- TGAAAATGGATGCCGACAG, tra RP- CTCTTTGGCGCAATCTTCTC, for dsx female transcript dsxFFP-CTATCCTTGGGAGCTGATGC, dsxF RP- TCGGGGCAAAGTAGTATTCG, for dsx male transcript dsxM FP- CAGACGCCAACATTGAAGAG, dsxM RP- CTGGAGTCGGTGGACAAATC, for msl2 FP- GTCACACTGGCTTCGCTCAG and msl2 RP- CCTGGGCTAGTTACCTGCAA were used.

### Validation of splicing results from time2splice using qRT-PCR and RT-PCR assays

Total RNA was extracted from fifty 0-2 Hr and 2-4 Hr female and male embryos expressing *MTD-GAL4>GFPRNAi* (con) and *MTD-GAL4>CLAMPRNAi* (CLAMP depleted). Sexed embryos were obtained as described in Reider et al 2017. We reverse-transcribed one microgram of total RNA using the SuperScript VILO cDNA Synthesis Kit (Life Technologies, USA) following the manufacturer’s protocol. We amplified target sequences by PCR using primers designed to span alternatively spliced junctions (**Fig S4**) listed in **Table S11** and Quick load Taq 2X Master mix (#M0271L, NEB, USA) according to the manufacturer’s protocol (28 cycles). 10ul of PCR product of each replicate for each gene was loaded in separate wells in 2% agarose gels and imaged using a ChemiDoc^TM^ MP Imaging system (BioRad, USA). All replicates for each gene were loaded on the same gel. The gel images were inverted and then quantified using the densitometry steps with the Fiji image analysis tool. qRT-PCR was carried out using 2X Azura Quant Green (#AZ-2120, Azura genomics, USA) according to the manufacturer’s instructions. Fold change between samples for each transcript was calculated the ýCT method (Schmittgen and Livak 2008). Student’s t-tests were performed to determine significant difference between groups (two samples at a time). Three replicates for qRT-PCR samples and four replicates for RT-PCR samples were performed.

### Immunoprecipitation

#### Nuclear and Cytoplasmic extract preparation

Male (S2) and female (Kc) cells were grown to a cell concentration of 2 x 10^6^ cells/mL in T25 tissue culture flasks. Cells were scraped from the flask, centrifuged for 5min at 2500rpm at 4°C. Supernatant was removed and cell pellets were washed twice in 5ml of cold PBS. The washed cell pellets were then resuspended in 5X volume of Buffer A (10mM HEPES pH7.9, 1.5mM MgCl_2_, 10mM KCl, 0.5mMDTT, 1X Protease inhibitors). Cells were incubated on ice for 15 minutes before dounce homogenization with an A pestle. Cytoplasmic fraction was collected after centrifugation at 4°C for 20 min at 700xg. The remaining nuclear pellet was re-suspended in 3 times volume in Buffer B (20 mM HEPES pH 7.9, 20% Glycerol, 0.5% NP 40, 200mMKCl, 0.5 mMEDTA, 1m MEGTA, 1X protease inhibitors). Nuclei after re-suspension were dounce homogenized with a B pestle. Nuclear debris was then pelleted by centrifugation at 10,000xg for 10 min at 4°C. 1 ml aliquots of both cytoplasmic and nuclear fractions were prepared in 1.5mL Protein LoBind Eppendorf tubes and flash frozen in liquid nitrogen for storage at −80 °C.

#### Immunoprecipitation

Magnetic anti-CLAMP beads were prepared to a final concentration of 10mg/mL by coupling rabbit anti-CLAMP antibody (SDIX) to magnetic beads, according to Dynabeads Antibody coupling kit (ThermoFisher Scientific) instructions. Similarly, Magnetic anti-FMRP beads were prepared using mouse anti-FMRP (5B6, DSHB, USA). Prepared anti-CLAMP, anti-FMRP and purchased anti-IgG (anti-rabbit IgG M-280 and anti-mouse IgG M-280 Dynabeads raised in sheep, Invitrogen, USA) were blocked to reduce background the night before the immunoprecipitation. First, the beads were washed 3 times for 5 minutes in 500L Tris-NaCl Wash (50mM Tris, 500mM NaCl, 0.1% NP-40) by rotating at 4C. The beads were next suspended in block buffer (3.3mg/mL of yeast tRNA extract prepared in 20mM HEPES, pH 7.9, 20% Glycerol, 0.5% NP-40, 200mM KCl, 1mM EDTA, and 2mM EGTA) and rotated overnight at 4C. The next day, beads were washed 3 times for 5 minutes in the block buffer without yeast tRNA by rotating at 4°C. After the final wash, beads were resuspended in the same amount of block buffer as the starting volume.

To 1mL of previously prepared nuclear extract, 100uL of blocked anti-CLAMP, anti-FMRP or anti-IgG magnetic Dynabeads were added. The nuclear extracts/cytoplasmic extracts and beads were then rotated for 1 hour at 4°C. Afterward, the beads were collected and the supernatant discarded. The beads were then washed three times in Tris-NaCl wash (50mM Tris, 500mM NaCl, 0.1% NP-40) by rotating for 5 minutes at 4°C and cleared by using a magnetic rack. To elute proteins from the beads, 100uL of 1% SDS was added, and the beads were boiled for 10 minutes at 95C. To the eluate, 300uL of ultrapure water was added, and the tubes gently vortexed. After collecting the beads on a magnetic rack, the eluate was saved in a clean Protein LoBind Eppendorf tube.

#### Western blotting

Squid and Hrb27C were detected in IP-CLAMP and IgG-rabbit protein samples using mouse anti-Squid (1:500, 1B11, DSHB) and rabbit anti-Hrb27C (1:5000, Fatima Gebauer), performed as mentioned above under western blotting protocol. CLAMP was detected in IP-FMRP and IgG-mouse samples using rabbit anti-CLAMP (1:1000).

### CUT&RUN

*CUT&RUN in embryos:*0-2 hr and 2-4 hr male and female embryos of desired genotypes (∼50 each) were collected on standard grape juice agar medium and washed with water. The embryos were dechorionated in 6% bleaching solution for 2 min and washed twice in ice cold 1XPBS. Centrifuged at 12,000g for 10 min at 4°C. Supernatants were discarded and embryos resuspended in 200μl Dig-Wash buffer with EDTA (20mM HEPES-NaOH, 150mM NaCl, 2mM EDTA, 0.5mM Spermidine, 10mM PMSF, 0.05% digitonin) and washed twice. Embryos were incubated in 200μl primary antibody overnight at 4°C on a tube rotator. Next, embryos were centrifuged at 12,000g for 10 min at 4°C and liquid removed and embryos were washed twice in Dig-Wash buffer with EDTA. Then, embryos were incubated for 3 hours at 4°C in ∼700 ng/ml pAMNase solution in Dig-Wash buffer with EDTA. Embryos were washed twice in Dig-Wash buffer without EDTA and resuspended in 150μl of Dig-Wash buffer without EDTA. Samples were equilibrated to 0°C on a heat block maintained on ice-bath. 2μl of 100mm CaCl_2_ added to each sample to initiate MNase activity and digestion was performed for 30 min before adding 150μl of 2X RSTOP Buffer (200mM NaCl, 20mM EDTA, 4mM EGTA, 50ug/ml RNase, 40ug/ml glycogen, 10pg/ml yeast spike-in DNA) to stop the reaction. Samples were tncubated at 37°C for 10 minutes to release the DNA fragments. Samples were spun at 12,000g for 10 minutes and aqueous layer transferred to a fresh 1.5 ml microfuge tube and centrifuged at 16,000g for 5 minutes. Cleared liquid was again transferred to a fresh tube, 1μl of 20% SDS and 2.5μl proteinase K (20ng/ml) added, incubated at 70°C for 10 minutes. 300μl PCI was added to each tube, mixed and total solution was transferred to phase lock tubes and centrifuged at 16,000g for 5 minutes. After adding 300μl of chloroform and mixing gently, samples were centrifuged at 16,000g for 5 minutes at RT. The aqueous layer was transferred to a DNA low binding tube. 1μl glycogen (5mg/ml) and 750μl ethanol added to precipitate DNA at −80°C. Samples were centrifuged at 16,000g for 10 min at 4°C and washed in ethanol twice. Pellet air dried and dissolved in 15μl of 1mM TrisHCl + 0.1mM EDTA pH 8.0 ^46, 47^. 1ng of Cut and Run DNA was used to make libraries using the KAPA Hyper prep kit and SeqCap adapters A &B (Roche) according to manufacturer’s protocol. For library amplification, 14 cycles were used and a 1.0X SPRI bead cleanup was performed using Agencourt Ampure XP beads. The following antibody concentrations were used: rabbit anti-CLAMP (5μg/sample, SDIX); 1:200 anti-rabbit (MilliporeSigma); rat anti-MLE (1:50, 6E11); 700ng/ml pA-MNase (from Steven Henikoff).

*CUT&RUN in cell lines*: Cells were allowed to grow to confluency and harvested. Equal number of cells for each category suspended in wash buffer and subjected to Cut&Run assay according to Skene et al 2018^47^ using rabbit anti-CLAMP (5µg) to immunoprecipitate CLAMP bound DNA fragments from male (S2) and female (Kc) cell lines. 3 replicates each for males and females were run, but during later stages one female sample was dropped due to insufficient starting material. Rabbit IgG was used as control, one for each male and female cell line sample. 1ng CUT&RUN DNA was used to generate libraries using Kapa Hyper prep kit (Roche, USA) and SeqCapAdapter Kit A (Roche, USA). 14 PCR cycles were used to amplify the libraries. AMPure XP beads (Beckman Coulter, USA) were used for library purification and fragment analysis was performed to check quality of the libraries made. Paired end 2×25 bp Illumina Hi-seq sequencing performed.

### RNA-sequencing

*RNA-seq in cell lines:* 15ug each of *clamp* dsRNA and GFP dsRNA used for *clamp* RNAi and GFPRNAi (con), respectively per T25 flask. Cells (Kc and S2) incubated with dsRNA in FBS minus media for 45 minutes and allowed to grow in media supplemented with 10% FBS for 6 days before harvesting. dsRNA targeting *gfp* (control) and *clamp* for RNAi have been previously validated and described^39, 91^. PCR products were used as a template to generate dsRNA using theT7 Megascript kit (Ambion, Inc., USA), followed by purification with the Qiagen RNeasy kit (Qiagen, USA). RNA was harvested using Rneasy mini plus kit (Qiagen, USA). 2 ug of total RNA was used for the construction of sequencing libraries. RNA libraries for RNA-seq were prepared using Illumina TruSeq V2 mRNA-Seq Library Prep Kit following the manufacturer’s protocols. Hi-seq paired end 100bp mRNA sequencing performed. Data was submitted to the GEO repository (#GSE220439). For gene expression analysis, the DESeq2 pipeline was used. For identifying CLAMP dependent splicing, our new time2splice pipeline was used.

*RNA-seq in third instar larvae (L3):* Total RNA was extracted from control (*yw*) and *clamp* mutant (*yw, clamp*^2^) male and female third instar larvae (3 each) using Trizol (Invitrogen, USA). Messenger RNA was purified from total RNA using poly-T oligo-attached magnetic beads. After fragmentation, the first strand cDNA was synthesized using random hexamer primers followed by the second strand cDNA synthesis. The library was ready after end repair, A-tailing, adapter ligation, size selection, amplification, and purification followed by paired-end RNA-sequencing in Illumina Novaseq 6000. The sequencing data was run through a SUPPA-based time2splice pipeline to identify CLAMP-dependent sex-specific splicing events. Data was submitted to the GEO repository (#GSE220455).

### iCLIP

Cells were allowed to grow to confluency and UV crosslinked using 254nm UV light in Stratalinker 2400 on ice (Stratagene, USA). UV treated cells were lysed to get different cellular fractions (Cytoplasmic, Nucleoplasmic and Chromatin) according to Fr-iCLIP (fractionation-iCLIP) protocol from Brugiolo et al 2017^42^. Chromatin and Nucleoplasmic fractions were sonicated with a Branson digital sonicator at 30% amplitude for 30 s total (10 sec on and 20 sec off) to disrupt DNA before IP. All three fractions were separately centrifuged at 20,000 xg for 5 min at 4℃. Fractions were tested by Western blotting using RNApolI for Chromatin Fraction, Actin for Cytoplasmic Fraction. Protein quantification for each fraction was done using manufacturer’s protocol for Pierce 660nm protein assay reagent (Thermo Scientific, USA). Each Fraction was subjected to iCLIP protocol as described in Huppertz et al 2014^41^ using rabbit-CLAMP antibody to immuniprecipitate bound RNAs which were extracted using proteinase K and phenol:chloroform. Custom cDNA libraries prepared according to Huppertz et al 2014^41^ using distinct primers Rt1clip-Rt16clip for separate samples containing individual 4nt-barcode sequences that allow multiplexing of samples. cDNA libraries for each sample amplified separately using 31 cycles of PCR, mixed together later and sequenced using standard illumina protocols. Heyl et al. 2020^92^ methods using the Galaxy CLIP-Explorer were followed to preprocess, perform quality control, post-process and perform peak calling.

## Computational Methods

### Time2splice tool

Time2splice is a new pipeline to identify temporal and sex-specific alternative splicing from multi-omics data that relies on the existing validated SUPPA method to identify differentially spliced isoforms (Trincado et al 2018). This pipeline combines SUPPA with several additional scripts to identify sex-specifically spliced genes and sex-biased genes at different time points.

Importantly, these scripts are partitioned into separate script files to enable the user to use only the scripts that they need for their analysis. Figure S1 describes the published methods and new scripts which we used in our analysis. Where boxes are numbered, the output from each step can be used as input for the subsequent step. **Step D** can be performed in any order depending on user needs. You can also see the README here (https://github.com/ashleymaeconard/time2splice) for a detailed description of the methods.

### a. Tutorial section for time2splice

Preprocess (scripts/preprocess): Retrieve raw data, quality control, trimming, alignment. Perform steps as needed.

`1_parse_sraRunTable.sh`

Creates `time2splice/` folder structure, as well as `metadatafile.csv` and `SraAccList.txt` (which is needed for next command to get .fastq files).

`1_get_fastq_files.sh`

Retrieves .fastq files by passing in `SraAccList.txt` from aforementioned step.

`2_run_fastQC.sh`

Runs FastQC for all .fastq files in a given directory.

`3_run_trim_galore.sh`

Run Trim Galore! followed by FastQC to trim any reads below quality threshold.

`3_merge_lines.sh`

Merges all the different lanes of the same flow cell .fastq files.

`4_run_Bowtie2.sh` or `preprocess/4_run_BWA.sh or preprocess/4_run_HISAT2.sh.`

Runs one or more of these three aligners (Bowtie2, BWA, or HISAT2) on .fastq data in a given directory.

`5_plot_alignment.py`

Plot the alignments from either one or two different aligners (Bowtie2 or HISAT2).

Temporal expression analysis (scripts/rna)

`1_run_salmon.sh`

Run salmon to quantify transcript expression for treatment and control samples.

e.g. .`/1_run_salmon.sh /nbu/compbio/aconard/larschan_data/sexed_embryo/`

`/data/compbio/aconard/splicing/results/salmon_results_ncbi_trans/`

`/data/compbio/aconard/BDGP6/transcriptome_dir/pub/infphilo/hisat2/data/bdgp6_ tran/genome.fa 3 10 1 _001.fastq.gz`

`2_run_suppa.sh`

Run Suppa for treatment and control samples.

e.g. `./2_run_suppa.sh /data/compbio/aconard/splicing/results/salmon_results/`

`/data/compbio/aconard/splicing/results/suppa_results_ncbi_trans/`

`/data/compbio/aconard/BDGP6/transcriptome_dir/pub/infphilo/hisat2/data/bdgp6_ tran/genome.fa 20`

`3_suppa_formatting.py`

Converts NM_ gene names to flybase name, then merging outputs from run_suppa (NM_ gene names by 1 TPM value column for each replicate)

`4_suppa.sh`

Identifies various forms of differential splicing (e.g. using PSI and DTU)

`5_calc_total_alt_splicing_controls.py#`

Calculate and plot the proportions of alternative splicing (in pie chart) in control samples.

`6_calc_total_alt_differential_splicing.py`

Calculate and plot the proportions of alternative splicing (in pie chart) in treatment samples.

`7_get_bias_genes.py`

Retrieve male and female biased genes and create bed files for average profile plotting.

`8_plots_splicing.ipynb`

Plotting transcript expression using PSI and DTU measures.

`8_alt_plots_splicing.ipynb`

Alternative code base to plot transcript expression using PSI and DTU measures.

`9_plots_splicing_time.ipynb`

Plot alternative splicing genes within categories (all females, all males, females sex specific, male sex specific, female all rest, male all rest, female non-sex specific, male non-sex specific, female new sex specific, male new sex specific) over time.

Temporal protein-DNA analysis (scripts/protein_dna)

`1_run_picard_markduplicates.sh`

Run Picard’s MarkDuplicates in for all .sorted.bam files in a given directory.

`2_run_macs2.sh`

Runs MACS2 to call peaks for all .sorted.bam files in a given directory.

`3_run_macs2_fold_enrich.sh`

Generate signal track using MACS2 to profile transcription factor modification enrichment levels genome-wide.

Temporal multi-omics integration (scripts/multio_analysis)

Note, there is no order to these scripts. Each analysis / results exploration is independent. More analysis scripts to come.

`overlap_protein_DNA_peaks.sh`

Runs Intervene to view intersection of each narrowpeak file.

`histogram_peak_val_intensity.ipynb`

Plot peak intensity for a given narrow peak file.

`get_coord_run_meme.sh`

Get coordinates of bed file and run through MEME.

`alt_splicing_chi_squared.ipynb`

Perform chi-squared test on alternative splicing categories. Mutually Exclusive Exons (MXE) used in this example.

### b. Identification of sex-specifically splicing events

We quantified the amount of alternative splicing using an exon-centric approach to quantify individual splice junctions by measuring percent spliced in (PSI) for a particular exon using SUPPA within time2splice.

PSI= IR (included reads)/ IR+ER (excluded reads)

The difference in PSI values (ýPSI) between samples implies differential inclusion or exclusion of alternative exons among the two sample types. For example, a positive ýPSI of 0.8 for an exon skip event means the exon is included in 80% of transcripts in the sample whereas a negative ýPSI value implies reduced inclusion of the alternative exon. First, we determined significant differences in ýPSI values for splicing events between the control female and male samples in 0-2 hr embryo (**Fig S2A**) and 2-4 hr embryo (**Fig S2D**) samples to identify CLAMP-independent sex-specific splicing differences between males and females. We have included volcano plots to show how we defined significant differences with a p-value cutoff of p-value<0.05. Next, we determined the splicing events which are significantly affected by *clamp* RNAi in female and male samples (**Fig S2B-C, E-F**). Lastly, we compared the lists of CLAMP-independent to CLAMP-dependent sex-specific splicing events identify the following categories of splicing events: 1) Splicing events that differ between wild type males and wild type females and are also dependent on CLAMP; 2) CLAMP-dependent new sex-specific splicing events: Splicing events that were not different when comparing wild type males and wild type females but do show sex-specific differences in the absence of CLAMP (**Fig 2B,C and Table S1**).

### c. Sex-specific splicing event analysis

RNA sequencing data from Rieder et al 2017 (#GSE102922), Kc and S2 cell line and third instar larval data generated by us were analyzed using time2splice to determine sex-specifically splicing events. dmel-all-r6.29.gtf from BDGP6 in genomes ^93^ was used to map each transcript identifier (ID) to gene ID and symbol, for .bed creation data for the associated chromosome, transcription start site (TSS) and transcription end site (TES), and strand information were imported from Illumina (https://support.illumina.com/sequencing/sequencing_software/igenome.html). From the raw data after quality control i.e, FastQC^94^, Salmon^95^ was used to quantify transcript expression for treatment and control samples. Calculated transcripts per million (TPM) values from SUPPA^26^ were used for all four replicates of female and male controls at both time points (before and after MZT). Each sample was filtered to include transcripts where the mean value is less than or equal to 3 TPMs per gene. The number of transcripts included at various thresholds were plotted from 1 to 10 and the fraction of genes filtered out begins to plateau around threshold 3. The percent of spliced in (PSI) transcripts between females and males were compared at both 0-2 Hr (pre-MZT) and 2-4 Hr (post-MZT); Kc and S2 cells; and third instar larval stage, L3 (p-value of 0.05), thereby resulting in delta PSI values and p-values for each transcription in each experimental condition comparison. Given these resulting delta transcript PSI values, significantly alternatively splice genes (p-value 0.05) were found between females vs. males 0-2 Hr (pre-MZT) controls to show which genes are normally sex-specifically spliced pre-MZT. The same process was followed at 2-4 Hr (post-MZT), in cell lines and third instar larvae. To then determine the sex-specifically spliced genes, the female RNAi experiment compared with the control delta PSI gave the number of total alternative spliced transcripts pre-MZT, then considering those that are not shared with males, and are only expressed in females normally, this defined our sex specifically spliced set of genes for females pre-MZT. This process was also performed for males pre-MZT, for post-MZT sample; for S2 and Kc cell lines and for female and male L3.

### d. Gene ontology analysis

Gene ontology (GO) analysis was performed using the R tool Clusterprofiler (Wu et al 2021). Specifically, time2splice’s script enrichment analysis. r implements GO analysis given an input gene set as a .txt file with a new line delimiter between genes. Given this input, it is converted to a vector of genes. The enrich GO function will return the enrichment GO categories after FDR correction. The FDR correction used is Benjamini Hochberg to account for the expected proportion of false positives among the variables (i.e. genes) for which we expect a difference. This was chosen over other methods such as the common Bonferroni method, as the Bonferroni correction controls the familywise error rate, where we are interested to account for false discoveries. The actual over-representation test itself is implemented in enrich GO according to Yu et al 2015 where they calculate a p-value using the hypergeometric distribution (Boyle et al 2011) and then perform multiple hypothesis correction. Importantly, while there are many tools to perform GO analysis, Cluster profiler was chosen due to its superior visuals and ability to handle multiple-omics types. This thus enables diverse additional analyses to be integrated into time2splice in the future such as ATAC-seq.

### ChIP-seq: Data analysis

We used preprocessed ChIP-seq data from Rieder et al 2019 (#GSE133637), specifically the .bw and .broadPeak.gz files in our analysis using ChIPseeker ^96^ and deeptools ^97^. Specifically, when plotting the average profiles using deeptools, we achieved a baseline signal representing genome-wide binding taking into consideration the number of genes in other groups by the following procedure: of all genes that are on (no zero read-count genes), we sampled the number of the largest other group (to which we are comparing), and ran computeMatrix on that subset. This process was repeated 500 times and the resulting 500 matrices were averaged to produce a representative signal. For motif analysis MEME ^98^ suite was used.

### CUT&RUN: Data analysis

Sequenced reads were run through FASTQC^94^(fastqc replicate_R1_001.fastq.gz replicate_R2_001.fastq.gz) with default parameters to check the quality of raw sequence data and filter out any sequences flagged for poor quality. Sequences were trimmed and reassessed for quality using TrimGalore (https://github.com/FelixKrueger/TrimGalore/issues/25) and FastQC ^94^, respectively. All Illumina lanes of the same flow cell. fastq files were merged, and sequenced reads were then mapped to release 6 *Drosophila melanogaster* genome (dm6). We compared Bowtie2^99^, HISAT2^100^, and BWA^101^. We found the best alignment quality with BWA and thus used this method’s results downstream. Next, we performed conversion to bam and sorting (e.g. using: bowtie2 -x dm6_genome -1 replicate_R1_001.fastq.gz -2 replicate_R2_001.fastq.gz -S out.sam > stout.txt 2> alignment_info.txt; samtools view -bS out.sam > out.bam; rm -rf out.sam; samtools sort out.bam -o out.sorted.bam). We removed reads (using samtools) with a MAPQ less than 30 and any reads with PCR duplicate reads (identified using MarkDuplicates Picard -2.20.2). Peaks identified using MACS2^102^( macs2 callpeak -t out.sorted.bam -B -f BAM --nomodel --SPMR -- keep-dup all -g dm --trackline -n outname --cutoff-analysis --call-summits -p 0.01 --outdir outdir) and keep duplicates separate. To calculate fold-enrichment macs2 is run again (macs2 bdgcmp -t $treat -c $control -o $out.sorted.bam_FE.bdg -m FE 2> $ out.sorted.bam_FE.log; macs2 bdgcmp -t $treat -c $control -o $out.sorted.bam_logLR.bdg -m logLR -p 0.00001 2). For motif analysis MEME ^98^ suite was used. Data submitted in GEO repository (#GSE174781, #GSE220981 and #GSE220053).

### iCLIP: Data analysis

The method from Heyl et al. 2020^92^ using the Galaxy CLIP-Explorer were followed to preprocess, perform quality control, post-process and perform peak calling. For preprocessing, UMI-Tools was used, and then UMI-tools and Cutadapt were used for Adapter, Barcode and UMI-removal. Cutadapt (Galaxy version 3.5) was used for filtering with a custom adapter sequence AGATCGGAAGAGCGGTTCAGCAGGAATGCCGAGACCGATCTCGTATGCCGTCTTCT GCTTG. All other settings followed the Heyl et al 2020 Galaxy iCLIP-explorer workflow. UMI-Tools Extract (Galaxy Version 1.1.2+galaxy2) was then used with a barcode pattern of NNNXXXXNN. No unpaired reads were allowed. The barcode was on the 3’ end. Je-Demultiplex (Galaxy Version 1.2.1) was then used for demultiplexing. FastQC was used for quality control. Mapping was done by RNA STAR (Galaxy version 2.5.2b-2) using dm6. All settings were chosen based on the existing parameters from the iCLIP-explorer settings. We selected FALSE for the option to use end-to-end read alignments with no soft-clipping. bedtools used for Read-Filtering, and UMI-Tools (Galaxy version 0.5.3.0) for de-duplication. PEAKachu was used for Peak Calling to generate bed files. The PEAKachu settings were followed using the Galaxy CLIP-explorer workflow. The maximum insert size was set to 150, the minimum cluster expression fraction was set to 0.01, the minimum block overlap set to 0.5, the minimum block expression set to 0.1. The Mad Multiplier was set to 0.0, the Fold Change Threshold was set to 2.0, and the adjusted p-value threshold was set to 0.05. Peaks were annotated using RCAS^103^ (RNA Centric Annotation System), a R package using Rstudio. MEME Suite used for motif detection. RCAS was used for functional analysis of the transcriptomes isolated by iCLIP, such as transcript features. ShinyGO 0.76^104^ was used to perform Gene Ontology Analysis of the iCLIP data. Data submitted in GEO repository (#GSE205987).

### a. Integrating CUT&RUN and iCLIP data

A Python script was created that iterates through all of the DNA peak bed files for CLAMP DNA binding sites in Kc and S2 cell lines (CUT&RUN data, #GSE220053) as a reference, and tests for overlap with CLAMP-bound RNA peaks (each sequence is between 25-50bp in size) in the Kc and S2 (iCLIP data, (#GSE205987). The overlaps are categorized into four main categories based upon the location of the overlap: 1) completely overlapping (purple lines in frequency plot), 2) partially overlapping at the DNA peak start site (red lines in frequency plot); 3) partially overlapping at the DNA peak end site (blue lines in frequency plot) and 4) non-overlapping, i.e. when there is an overlap in a region outside the DNA binding site (yellow lines in frequency plot). This extended region is defined by the *scope* variable in the script, allowing the overlap to look for binding sites in the proximity of the DNA binding site (this scope is 2 kb including the DNA binding site). It should be noted that multiple RNA peaks can be found on one DNA peak. All of these overlaps are placed onto a [-scope, scope] region. Then, each type of overlap shown with a different color is overlaid and plotted onto a frequency plot. So, if the frequency at a given base pair is 5, then there are five overlaps that contained that base pair within the region defined by the scope.

### b. Identifying RBP motif in iCLIP data

The online tool cisBP-RNA database (cisbp-rna.ccbr.utoronto.ca) was used to identify binding motifs for six proteins in *Drosophila Melanogaster*, namely Sxl, Tra2, Hrb87F/hrp36, Hrb98DE/hrp38, Hrb27C, and Lark, within iCLIP data for CLAMP bound RNAs in S2 and Kc cell chromatin and nucleoplasmic fractions. First, the RNA Binding Protein (RBP) information for each protein was extracted from the database and placed in the cart using the search bar on the home page. Then, the RNA Scan tool was run for all the RBPs in the cart to scan for the RBP motifs in the list of CLAMP-bound RNA sequences, with the inputs of the FASTA sequence for each fraction, the species set to *Drosophila melanogaster*, and the motif model used was PWMs (Energy) with a threshold of 0.2. From there, the resulting CSV files were passed through a Python script to count the number of CLAMP binding motifs per fraction that contained each protein. Then, the frequency of binding sites for each RNA binding protein within each sample was plotted as a fraction on separate graphs.

## Acknowledgments

This work and funding to M.R. was supported by R35GM126994 to E.N.L. from NIH. A.M.C is funded by the NSF Graduate Research Fellowship and CCMB, Brown University. We thank Bloomington stock center for fly lines. We thank Leila Rieder for SD and Rsp^s^ stocks, Peter Becker for MSL2 and MLE antibodies, Steve Henikoff for pAMNase protein and spike-in DNA for Cut and Run, and Daniel J McKay for sharing his Cut and Run protocol for tissues.

## Author Contributions

**M.R**., **A.M.C**. and **E.N.L**. planned experiments, analyzed results and wrote the manuscript. **A.M.C** did all the computational analysis regarding time2splice pipeline. **M.R.** carried out the experimental work and collected data for Cut and Run, iCLIP, Polytene squashes and IF, splicing assays and IP. **J.U.** carried out mRNA-sequencing in cell lines, sex determination pathway splicing assays and WB. **J.A** analyzed the MLE cut and run data and performed mRNA-sequencing in L3. **A.H** analyzed the iCLIP-seq data. **P.M** analyzed the cell line and third instar larval RNA-seq data using time2splice pipeline to identify CLAMP dependent splicing events and integrated CLAMP iCLIP data with CLAMP CUT&RUN in cell lines. **S.V** did the motif analysis to identify motifs for RBPs involved in splicing in CLAMP iCLIP data.

## Figure legends

**Fig S1.**
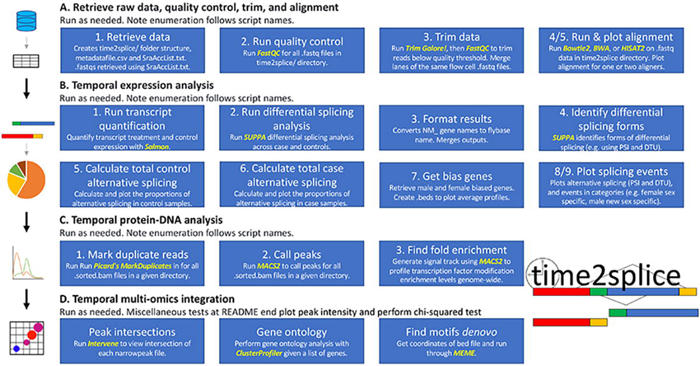
Schematic diagram describing each step-in sequential order performed by the time2splice pipeline.

**Fig S2.**
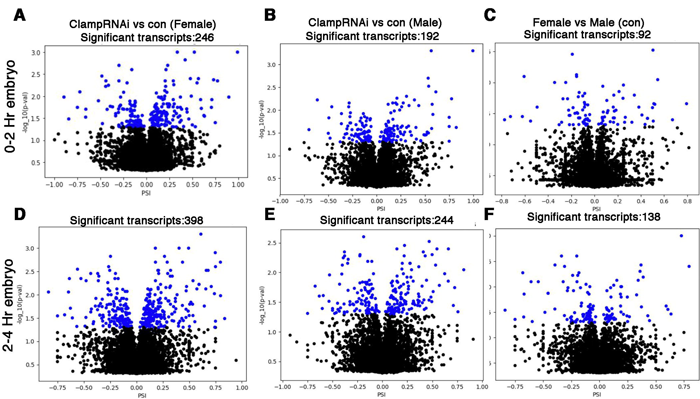
Sex-specific differences in alternative splicing in early *Drosophila melanogaster* embryos. **A-F** Volcano plots showing log_10__pvalues for significant differences between PSI values for splicing events at early embryonic stages in female and male embryos 0-2 Hr (**A, C, E**) and 2-4 Hr (B, **D, F**). Significant changes are labeled as blue dots (p<0.05 and PSI minimum ±0.2). For example, PSI of +0.8 means 80% of the transcripts retained the exon, while negative PSI values mean reduced inclusion of the alternative exon.

**Fig S3.**
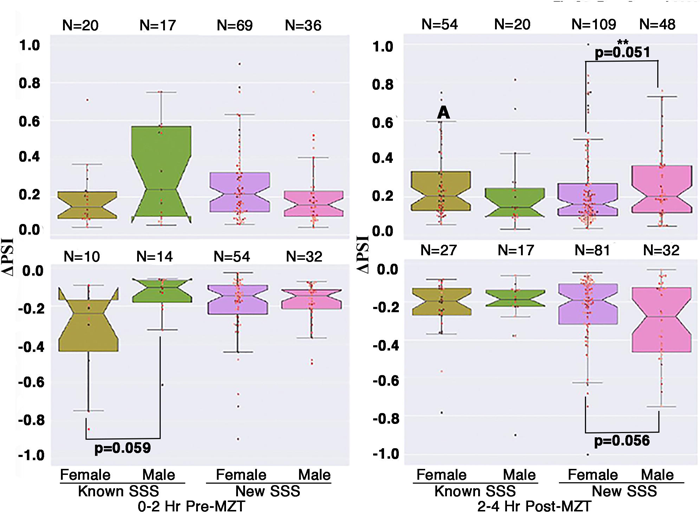
CLAMP inhibits aberrant alternative splicing in post-MZT male embryos. Box plot showing ΔPSI values for **known** (different between females and males in control samples) and **new** (not different in males and females in control samples) CLAMP-dependent sex-specific spliced events at 0-2Hr/pre-MZT and 2-4Hr/post-MZT female and male embryos. **N** denotes the total number of splicing events in each category, and p-values for groups showing significant differences are noted at the bottom of the line connecting the compared groups.

**Fig S4.**
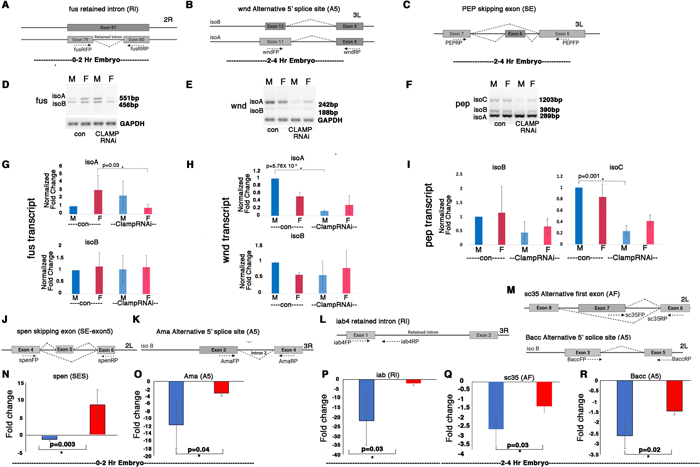
Validation of splicing differences at randomly chosen target genes where CLAMP regulates sex-specific splicing by RT-PCR and qRT-PCR. **A-C.** Schematic showing alternative splicing events resulting in different isoforms of the same gene which are regulated by CLAMP and the position of primers (dotted arrows) used to detect these isoforms in RT-PCR assays. **D-F**. Inverted agarose electrophoretic gel images show the expression level of each isoform detected using primers in the RT-PCR assays noted in **A-C** in male (**M**) and female (**F**) early embryos under control *GFPRNAi* as well as *CLAMPRNAi* conditions. **G-I.** Bar plots showing the change in levels of specific isoforms resulting from alternative splicing events in male (**blue**) and female (**red**) early embryos under control *GFPRNAi* (deeper shade of blue and red) and *CLAMPRNAi* (lighter shade of blue and red) conditions. The isoform transcript levels are normalized by the levels of ***gapdh*** housekeeping gene transcript. p-values (paired student’s t-test) for groups showing significant differences (*) are noted at the top of the line connecting the compared groups (**four replicates for each gene**). **J-M.** Schematic showing alternative splicing events resulting in different isoforms of the same gene which are regulated by CLAMP and the position of primers (dotted arrows) used to detect these isoforms by qRT-PCR analysis. **N-R.** Bar plot showing fold changes in transcript levels of the isoform detected using primers shown in **J-M** of respective genes by qRT-PCR (**three replicates**) in the *MTD-GAL4>CLAMPRNAi* genotype when compared to the control (*MTD-GAL4>GFPRNAi*) genotype, in 0-2 Hr/pre-MZT and 2-4 Hr/post-MZT sexed embryos. Fold changes for each transcript differ significantly between males (**blue**) and females (**red**) (pý 0.05, Student t-test).

**Fig S5.**
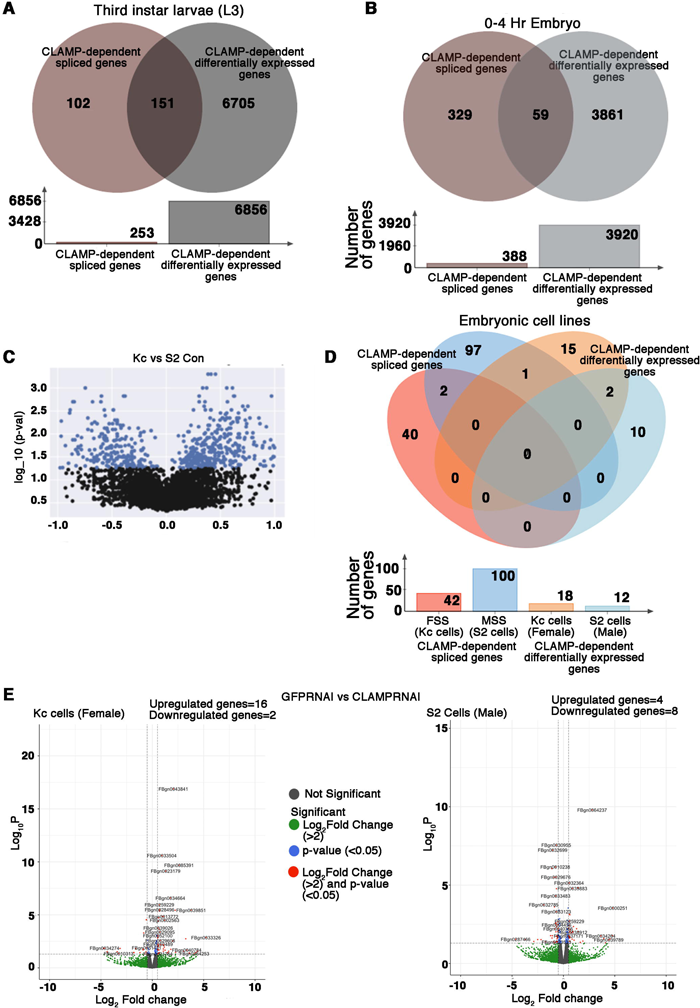
CLAMP has context specific dual role in splicing and transcription at specific genomic loci. **A-B.** Venn diagram showing overlap between CLAMP dependent spliced genes with CLAMP-dependent differentially expressed genes in third instar larvae (A) and 0-4 Hr Embryo (B). The total number of genes in each category is shown in the bar plot below the Venn diagram. **C.** Volcano plot showing log_10__p-values for significant differences between PSI values for splicing events in female (Kc) and male (S2) *Drosophila* embryonic cell lines. Significantly changed splicing events (**N=615**) are labeled as blue dots (p<0.05 and PSI minimum ±0.2). **D.** Venn diagram showing overlaps between dependent spliced genes in Kc (female) cells (pink circle) and S2 (male) cells (deep blue circle) with CLAMP dependent differentially expressed genes in Kc (orange circle) and S2 cell lines (light blue circle). Bar plot shows the total number of genes in each category. **E**. Volcano plots showing differential gene expression in Kc (female) and S2 (male) cell lines after *clamp* RNAi compared to control (*GFP* RNAi).

**Fig S6.**
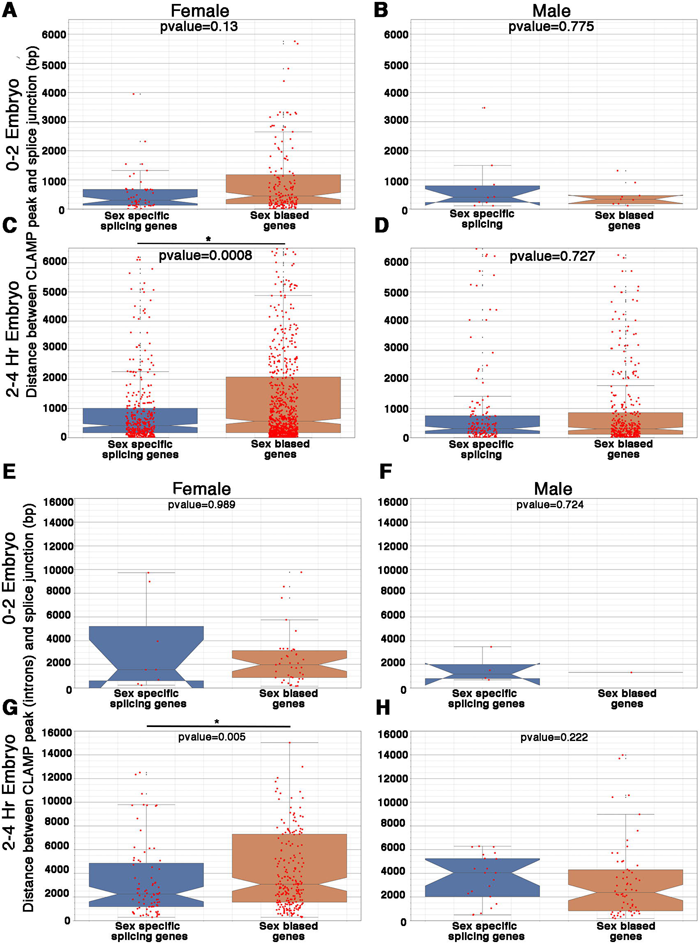
CLAMP binds to chromatin near splice junctions. **A-D.** Notched box plots representing distance between **CLAMP peaks** in sex-specifically spliced and sex-biased genes with the nearest splice junction in female (**A, C**) and male (**B, D**) 0-2 Hr (**A, B**) and 2-4 Hr (**C, D**) embryos. p-values (Mann-Whitney test) for each group are noted at the top and those with significant differences and the compared groups are connected with a solid black line with an asterisk at the top *. t-tests and KS-tests were also performed and showed the same results. **E-H.** Notched box plots representing distance between **CLAMP peaks in introns** of the sex-specifically spliced and sex-biased genes and the nearest splice junction in female (**E, G**) and male (**F, H**) 0-2 Hr (**E, F**) and 2-4 Hr (**G, H**) embryos. p-values (Mann-Whitney test) for each group are noted at the top for those with significant differences and the compared groups are connected with a solid black line with an asterisk at the top *. t-test and KS-test were also performed and showed the same results.

**Fig S7.**
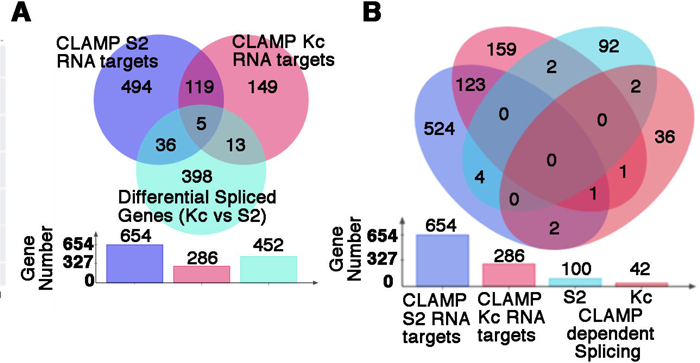
CLAMP regulates splicing both indirectly as well as directly in *Drosophila* embryonic cell lines. **A.** Venn diagram showing overlaps between CLAMP RNA targets in S2 (male) cells (violet circle) and Kc (female) cells (deep pink circle) with genes differentially spliced (**N=452**) between Kc and S2 cell lines (green circle). Bar plot shows the total number of genes in each category. **B.** Venn diagram showing overlap between CLAMP iCLIP RNA targets in S2 (male) cells (violet circle) and Kc (female) cells (deep pink circle) with CLAMP-dependent spliced genes in S2 (turquoise blue) and Kc (light pink circle) cells. The total number of genes in each category is shown in the bar plot below the Venn diagram.

**Fig S8.**
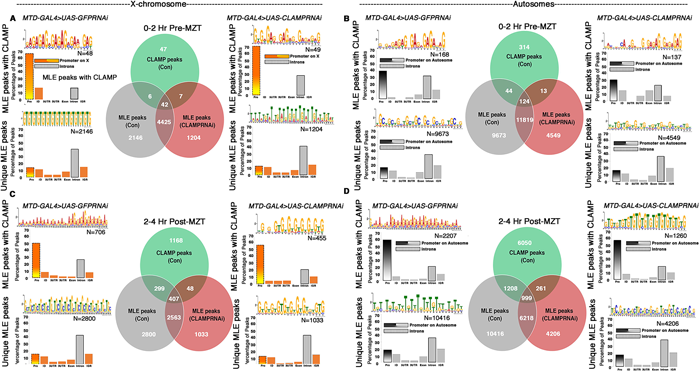
MLE binds to different motifs and to different chromatin regions when colocalizing with CLAMP compared to its unique binding sites that lack CLAMP. **A-D** Bar plots show the distribution of MLE peak percent overlap with CLAMP peaks and unique MLE peaks at different types of genomic regions: 1) **Pro**=Promoter; 2) **ID**=Immediate downstream; 3) **5UTR**=5’ untranslated region; 4) **3UTR**= 3’ untranslated region; 5) **Exon**; 6) **Intron**; and 7) **IGR**=Intergenic region. MLE distribution was measured on the X-chromosome (**A, C**) and autosomes (**B, D)** in male embryos at 0-2Hr/pre-MZT (**A, B**) and 2-4Hr/post-MZT (**C, D**) stages under normal conditions and after the loss of maternal CLAMP. ‘**N**’ denotes the total number of peaks in each category. The most frequently identified sequence motif (MEME) for MLE peaks overlapping with CLAMP and unique MLE peaks in each category is shown at the top of the relevant bar plot. Venn diagrams in **A-D** compare CLAMP peaks in control (green circle) with MLE peaks in control (grey circle) to determine the overlapping peaks (intersection between the two) and unique MLE peaks. The red circle denotes MLE peaks that are present in absence of maternal CLAMP. The MLE peaks lost in absence of maternal CLAMP (exclusively grey area) and gained (exclusively red area).

**Fig S9.**
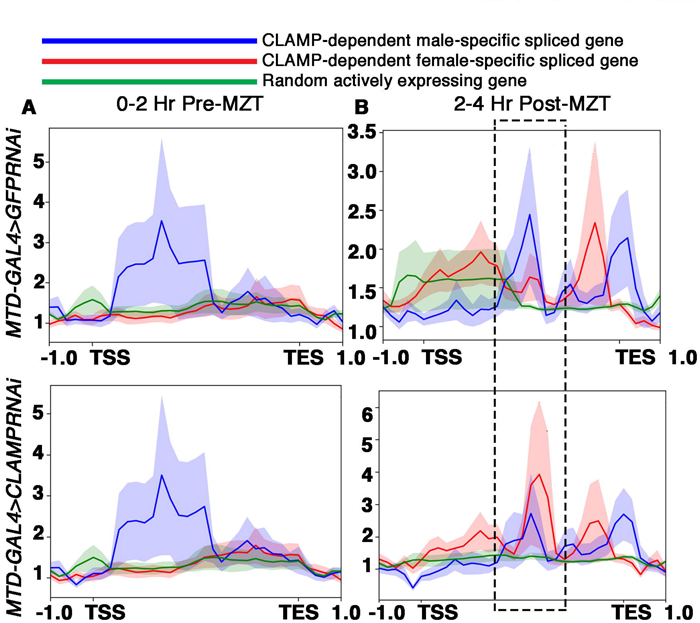
MLE binding at CLAMP-dependent sex-specifically spliced genes. A-B. Average profile for MLE over the gene bodies of CLAMP-dependent male (blue line) and female (red line) sex specifically spliced genes in 0-2 Hr pre-MZT (**G**) and 2-4 Hr post-MZT (**H**) male embryos under normal condition (**top**) and after the loss of maternal CLAMP (**bottom**). A set of random active genes were used as a control (**green line**). **TSS**=**T**ranscription **s**tart **s**ite and **TES**= **T**ranscription **e**nd **s**ite. The rectangular box with dashed lines in panel **H** shows that *CLAMP* RNAi increases binding of MLE to female sex-specifically spliced genes (**red line**) compared to male sex-specifically spliced genes (**blue line**) compared to control RNAi.

**Fig S10.**
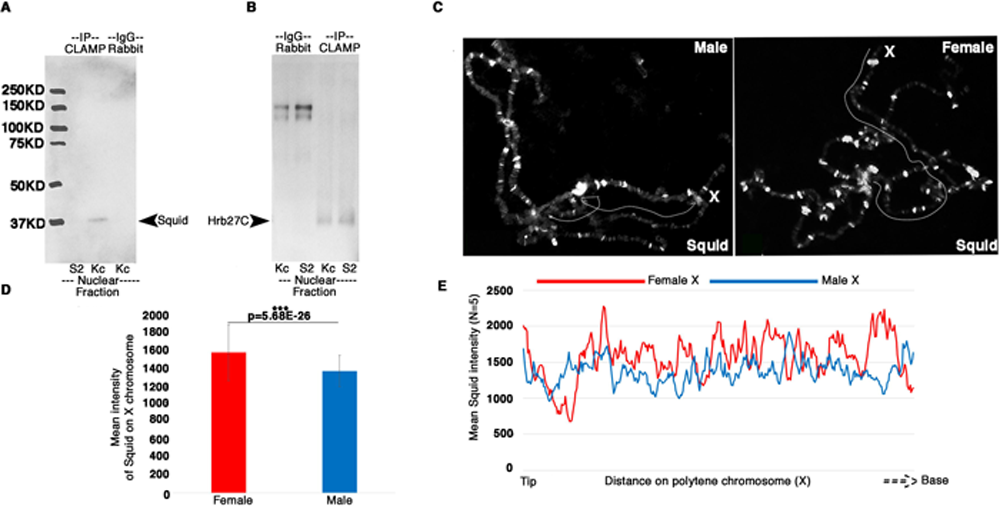
CLAMP sex-specifically interacts with components of the spliceosome complex. **A-B**. Western blot for Squid (hrp40, **A**) and Hrb27C (hrp48, **B**) in nuclear protein fractions from Kc (female) and S2 (male) cells subjected to IP (Immunoprecipitation) using rabbit anti-CLAMP. Rabbit IgG was used as control (lane 4, **A**, and lanes 1&2, **B**). **C.** Fluorescent microscopy images show the distribution of Squid (white) on chromatin (grey) in polytene chromosome preparations from third instar larval salivary glands. The dotted white line indicates the X-chromosome. **D.** Bar plot showing the average intensity of Squid immunostaining on female (red) and male (blue) X-chromosomes. Intensities of Squid immunostaining at different genomic locations of the X-chromosome were summed and divided by number of genomic locations to obtain an average intensity (data obtained using plot profile in Fiji) (**N=5**) **E.** The mean intensity profile for Squid along a portion of the female (red) and male (blue) X-chromosome polytene spread (**N=5**).

**Fig S11.**
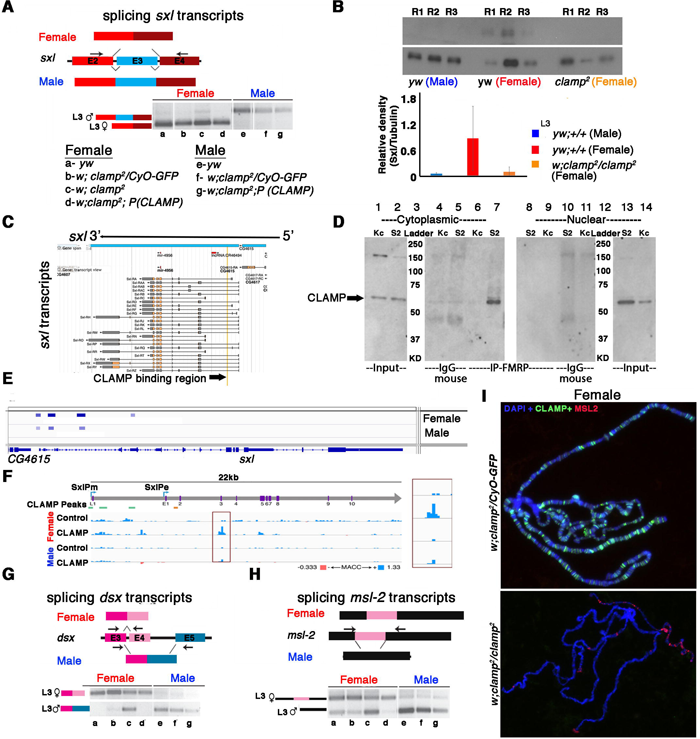
Alternative splicing of components of the sex determination pathway is regulated by zygotic CLAMP in females. **A.** Electrophoresis gel image (inverted colors) showing splicing of *sxl* transcripts in third instar larvae of females and males of genotypes listed in the key (a-g) with a representative schematic at the top of the gel image. **B.** Western blot showing the level of Sxl protein in genotypes (3 replicates for each) mentioned below each lane. Tubulin levels were used as a protein loading control. Below the blot is the relative quantification of Sxl protein levels compared with Tubulin and each genotype is represented by separately colored bars. **C.** BLAST alignment showing CLAMP binding at the 5’UTR regions of *sxl* transcripts from iCLIP data. **D.** Western blot for CLAMP in cytoplasmic and nuclear protein fractions from Kc (female) and S2 (male) cells after IP (immunoprecipitation) using mouse anti-FMRP. IgG-mouse was used as negative control (lanes 4, 5 and lanes 11 & 12). **E** IGV browser screenshot showing CLAMP peaks (rectangular boxes in light blue) at the genomic locus for the *sxl* gene in male and female cell lines. For each category, the narrow peak file is shown. **F.** Chromatin accessibility measured by the MNase Accessibility (MACC) score is shown across the *sxl* gene in male (S2) and female (Kc) cells under control and *clamp* RNAi conditions. The MACC score is a previously reported (Urban et al. 2017) quantification of chromatin accessibility at each locus in the genome. Positive accessibility values (blue) indicate high chromatin accessibility, and negative (red) accessibility values indicate low chromatin accessibility. Each window covers MACC values ranging from -0.333 to +1.33. MACC values increase in females after *clamp* RNAi, specifically at exon 3 (red box), and are shown in the inset to the right. Green boxes represent CLAMP binding peaks in the *sxl* gene just below the schematic for the *sxl* gene itself. **G-H.** Electrophoresis gel image from third instar larval samples (a-g) showing splicing of *dsx* (**G**) and *msl2* (**H**) transcripts in females (lane sa-d) and males (lanes e-g). a-g genotypes are the same as in panel **A**. The schematics at the top of each gel image show female and male splice variants of *dsx* (**G)** and *msl2* (**H**) transcripts. **I.** Fluorescent microscopy images of polytene chromosomes from the third instar salivary gland in the genotypes listed to the left of each panel (heterozygous control and *clamp*^2^ null) show the distribution of CLAMP (green) and MSL2 (red) on chromatin (blue, DAPI).

**Fig S12.**
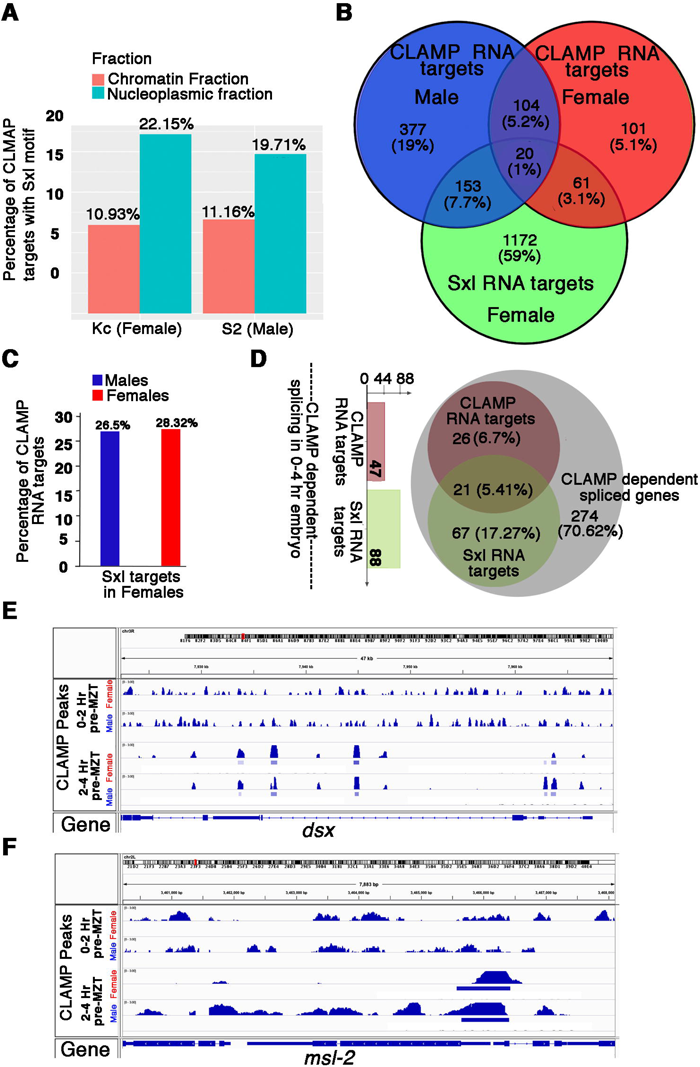
CLAMP and Sxl have common and independent RNA targets. **A.** Bar plots showing the percentage of CLAMP RNA targets that contain the Sxl RNA-binding motif in male and female nuclear fractions. **B**. Venn diagrams comparing CLAMP RNA targets between male and female *Drosophila* cell lines and between Sxl RNA targets identified previously in the adult*Drosophila* female head. **C**. Bar plots showing the percentage of CLAMP male **(blue)** and female (**red**) iCLIP targets that are also Sxl iCLIP targets. **D.** Venn diagram comparing CLAMP-dependent spliced genes in early embryos (0-4 Hr) with iCLIP RNA targets of CLAMP and Sxl. The total number of CLAMP-dependent spliced genes in early embryos which are also direct iCLIP RNA targets for CLAMP (red) and Sxl (green) is shown in the corresponding bar plots. **E-F** IGV browser screen shot showing CLAMP peaks (rectangular boxes in light blue) at the genomic locus for the ***dsx*** (**E**) and ***msl-2*** (**F**) genes in male and female 0-2 Hr/pre-MZT and 2-4Hr/ post-MZT embryos. The bigwig file (upper track) and the corresponding narrow peak file (lower track) are both shown.

**Fig S13.**
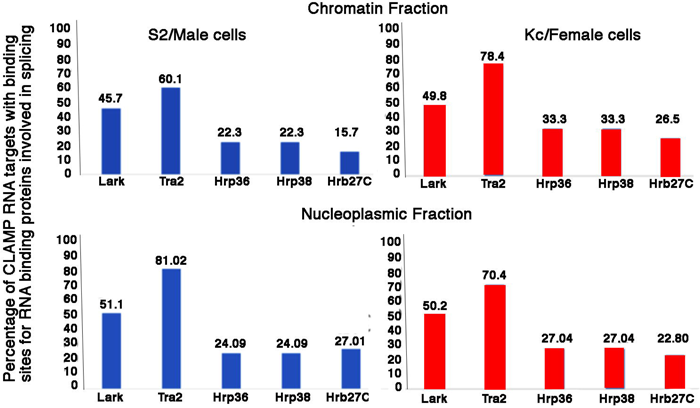
Multiple RNA binding proteins involved in splicing have target motifs in CLAMP bound RNA. Bar plots showing the percentage of iCLIP CLAMP RNA targets in the S2 cell chromatin fraction (**N=645**), S2 cell nucleoplasmic fraction (**N=53**), Kc cell chromatin fraction (**N=203**) and Kc cell nucleoplasmic fraction (**N=119**) which have RNA binding sequence motifs for other RNA binding proteins involved in alternative splicing (noted along the x-axis).

## Table legends

**Table S1:** List of CLAMP dependent sex-specific splicing events

**Table S3:** List of all and sex-specific splicing events regulated by CLAMP in *Drosophila* third instar larvae (L3) and which of them are direct CLAMP targets.

**Table S4:** List of all CLAMP dependent differentially expressed genes in Drosophila male and female third instar larvae (L3)

**Table S5:** List of a) Differential splicing events between Kc (female) and S2 (male) cell lines and which of them are direct CLAMP RNA targets, b) All CLAMP dependent splicing events in *Drosophila* sexed embryonic cell lines, c) CLAMP dependent female and male specific-splicing events in *Drosophila* sexed embryonic cell lines and which of them are direct CLAMP targets.

**Table S6:** List of CLAMP dependent spliced genes which are directly bound by CLAMP, list of sex-specifically spliced genes and sex-biasedly expressed genes in 0-2 and 2-4 Hr male and female Pre-MZT and Post-MZT embryos, respectively.

**Table S7:** CLAMP RNA targets identified in the nuclear fractions of male (S2) and female (Kc) cells. List of CLAMP dependent spliced genes which are direct CLAMP RNA targets.

**Table S8:** List of common RNA targets of CLAMP and SXL.

**Table S9:** List of CLAMP dependent male and female specifically spliced genes which are direct SXL targets.

**Table S10:** List of CLAMP dependent male and female specifically spliced genes which are common and unique direct targets of CLAMP and SXL.

**Table S2:**
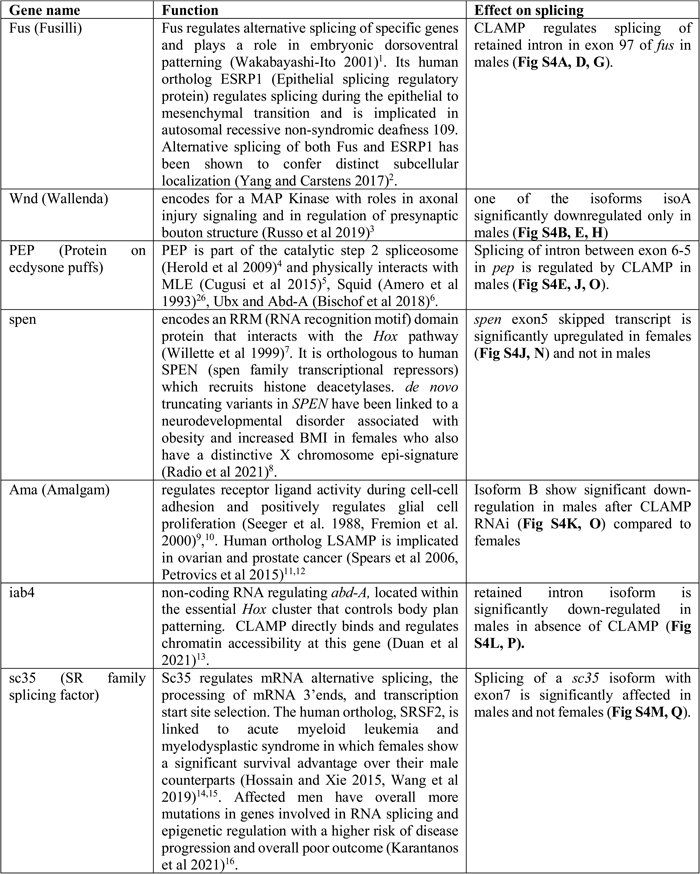
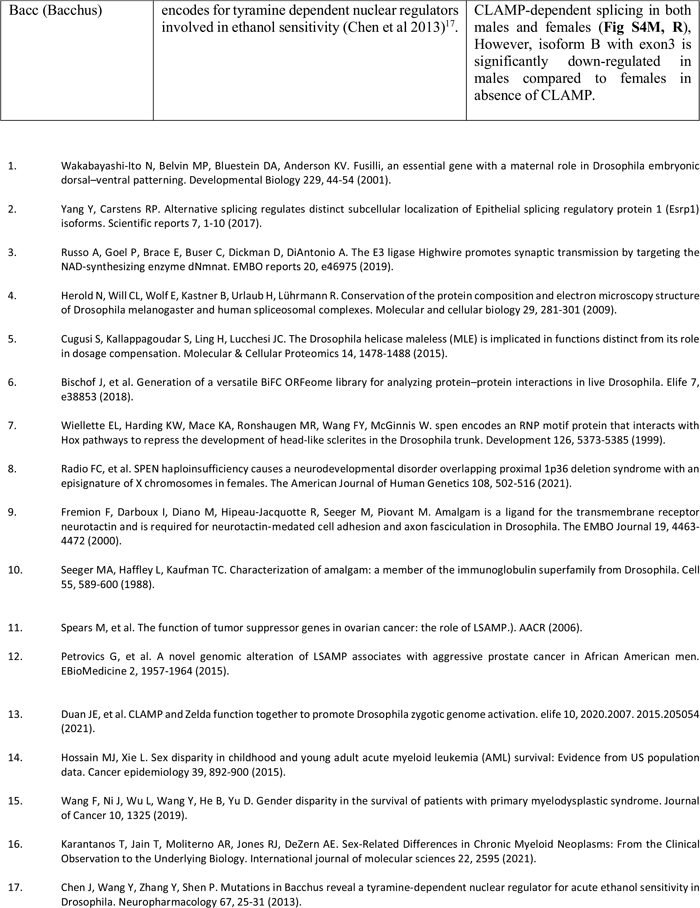
Summarizing the results and functions of the validated target genes at which splicing is regulated by CLAMP.

**Table S11:**
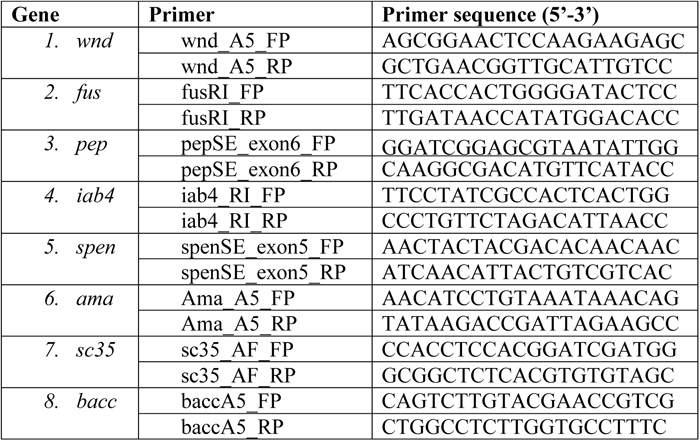
List of primers used for validation.

## References

1. Horn, T., Gosliga, A., Li, C., Enculescu, M. & Legewie, S. Position-dependent effects of RNA-binding proteins in the context of co-transcriptional splicing. npj Systems Biology and Applications 9, 1 (2023).

2. Shukla, S. & Oberdoerffer, S. Co-transcriptional regulation of alternative pre-mRNA splicing. Biochimica et Biophysica Acta (BBA)-Gene Regulatory Mechanisms 1819, 673–683 (2012).

3. Bedi, K. et al. Cotranscriptional splicing efficiencies differ within genes and between cell types. RNA 27, 829–840 (2021).

4. Gehring, N.H. & Roignant, J.-Y. Anything but ordinary–emerging splicing mechanisms in eukaryotic gene regulation. Trends in Genetics 37, 355–372 (2021).

5. Herzel, L., Ottoz, D.S., Alpert, T. & Neugebauer, K.M. Splicing and transcription touch base: co-transcriptional spliceosome assembly and function. Nature reviews Molecular cell biology 18, 637–650 (2017).

6. Merkhofer, E.C., Hu, P. & Johnson, T.L. Introduction to cotranscriptional RNA splicing. Spliceosomal Pre-mRNA Splicing: Methods and Protocols, 83–96 (2014).

7. Zhang, S. et al. Structure of a transcribing RNA polymerase II–U1 snRNP complex. Science 371, 305–309 (2021).

8. Fu, X.-D. & Ares Jr, M. Context-dependent control of alternative splicing by RNA-binding proteins. Nature Reviews Genetics 15, 689–701 (2014).

9. Naftelberg, S., Schor, I.E., Ast, G. & Kornblihtt, A.R. Regulation of alternative splicing through coupling with transcription and chromatin structure. Annual review of biochemistry 84, 165–198 (2015).

10. Ji, X. et al. SR proteins collaborate with 7SK and promoter-associated nascent RNA to release paused polymerase. Cell 153, 855–868 (2013).

11. Van Nostrand, E.L. et al. A large-scale binding and functional map of human RNA-binding proteins. Nature 583, 711–719 (2020).

12. Brooks, A.N. et al. Regulation of alternative splicing in Drosophila by 56 RNA binding proteins. Genome research 25, 1771–1780 (2015).

13. Schulz, K.N. et al. Zelda is differentially required for chromatin accessibility, transcription factor binding, and gene expression in the early Drosophila embryo. Genome research 25, 1715–1726 (2015).

14. Schulz, K.N. & Harrison, M.M. Mechanisms regulating zygotic genome activation. Nature Reviews Genetics 20, 221–234 (2019).

15. Boumpas, P., Merabet, S. & Carnesecchi, J. Integrating transcription and splicing into cell fate: transcription factors on the block. Wiley Interdisciplinary Reviews: RNA, e1752 (2022).

16. Rieder, L.E. et al. Histone locus regulation by the Drosophila dosage compensation adaptor protein CLAMP. Genes & development 31, 1494–1508 (2017).

17. Artieri, C.G. & Fraser, H.B. Transcript length mediates developmental timing of gene expression across Drosophila. Molecular biology and evolution 31, 2879–2889 (2014).

18. De Renzis, S., Elemento, O., Tavazoie, S. & Wieschaus, E.F. Unmasking activation of the zygotic genome using chromosomal deletions in the Drosophila embryo. PLoS Biol 5, e117 (2007).

19. Guilgur, L.G. et al. Requirement for highly efficient pre-mRNA splicing during Drosophila early embryonic development. Elife 3, e02181 (2014).

20. Förch, P. & Valcárcel, J. Splicing regulation in Drosophila sex determination. In Regulation of Alternative Splicing 127–151 (Springer, 2003).

21. Urban, J. et al. Enhanced chromatin accessibility of the dosage compensated Drosophila male X-chromosome requires the CLAMP zinc finger protein. PloS one 12, e0186855 (2017).

22. Duan, J.E., et al. CLAMP and Zelda function together as pioneer transcription factors to promote Drosophila zygotic genome activation. bioRxiv (2020).

23. Urban, J.A., Urban, J.M., Kuzu, G. & Larschan, E.N. The Drosophila CLAMP protein associates with diverse proteins on chromatin. PloS one 12, e0189772 (2017).

24. Kaye, E.G. et al. Differential occupancy of two GA-binding proteins promotes targeting of the drosophila dosage compensation complex to the male X chromosome. Cell reports 22, 3227–3239 (2018).

25. Quinn, J.J. et al. Rapid evolutionary turnover underlies conserved lncRNA–genome interactions. Genes & development 30, 191–207 (2016).

26. Trincado, J.L. et al. SUPPA2: fast, accurate, and uncertainty-aware differential splicing analysis across multiple conditions. Genome biology 19, 1–11 (2018).

27. Urban, J.A. et al. The essential Drosophila CLAMP protein differentially regulates non-coding roX RNAs in male and females. Chromosome Research 25, 101–113 (2017).

28. Rieder, L.E., Jordan III, W.T. & Larschan, E.N. Targeting of the dosage-compensated male X-chromosome during early Drosophila development. Cell reports 29, 4268–4275. e2 (2019).

29. Duan, J.E. et al. CLAMP and Zelda function together to promote Drosophila zygotic genome activation. elife 10, 2020.07.15.205054 (2021).

30. Atallah, J. & Lott, S.E. Evolution of maternal and zygotic mRNA complements in the early Drosophila embryo. PLoS genetics 14, e1007838 (2018).

31. Kwasnieski, J.C., Orr-Weaver, T.L. & Bartel, D.P. Early genome activation in Drosophila is extensive with an initial tendency for aborted transcripts and retained introns. Genome research 29, 1188–1197 (2019).

32. Gutierrez-Perez, I. et al. Ecdysone-induced 3D chromatin reorganization involves active enhancers bound by Pipsqueak and Polycomb. Cell reports 28, 2715–2727.e5 (2019).

33. Kidder, B.L., Hu, G. & Zhao, K. ChIP-Seq: technical considerations for obtaining high-quality data. Nature immunology 12, 918–922 (2011).

34. Tsompana, M. & Buck, M.J. Chromatin accessibility: a window into the genome. Epigenetics & chromatin 7, 1–16 (2014).

35. Cherbas, L. & Gong, L. Cell lines. Methods 68, 74–81 (2014).

36. Cherbas, L., Moss, R. & Cherbas, P. Transformation techniques for Drosophila cell lines. Methods in cell biology 44, 161–179 (1994).

37. Alekseyenko, A.A. et al. A sequence motif within chromatin entry sites directs MSL establishment on the Drosophila X chromosome. Cell 134, 599–609 (2008).

38. Straub, T., Zabel, A., Gilfillan, G.D., Feller, C. & Becker, P.B. Different chromatin interfaces of the Drosophila dosage compensation complex revealed by high-shear ChIP-seq. Genome research 23, 473–485 (2013).

39. Hamada, F.N., Park, P.J., Gordadze, P.R. & Kuroda, M.I. Global regulation of X chromosomal genes by the MSL complex in Drosophila melanogaster. Genes & development 19, 2289–2294 (2005).

40. Soruco, M.M. et al. The CLAMP protein links the MSL complex to the X chromosome during Drosophila dosage compensation. Genes & development 27, 1551–1556 (2013).

41. Huppertz, I. et al. iCLIP: protein–RNA interactions at nucleotide resolution. Methods 65, 274–287 (2014).

42. Brugiolo, M., Botti, V., Liu, N., Müller-McNicoll, M. & Neugebauer, K.M. Fractionation iCLIP detects persistent SR protein binding to conserved, retained introns in chromatin, nucleoplasm and cytoplasm. Nucleic acids research 45, 10452–10465 (2017).

43. Harrison, A.F. & Shorter, J. RNA-binding proteins with prion-like domains in health and disease. Biochemical Journal 474, 1417–1438 (2017).

44. March, Z.M., King, O.D. & Shorter, J. Prion-like domains as epigenetic regulators, scaffolds for subcellular organization, and drivers of neurodegenerative disease. Brain research 1647, 9–18 (2016).

45. Blanchette, M. et al. Genome-wide analysis of alternative pre-mRNA splicing and RNA-binding specificities of the Drosophila hnRNP A/B family members. Molecular cell 33, 438–449 (2009).

46. Uyehara, C.M. & McKay, D.J. Direct and widespread role for the nuclear receptor EcR in mediating the response to ecdysone in Drosophila. Proceedings of the National Academy of Sciences 116, 9893–9902 (2019).

47. Skene, P.J., Henikoff, J.G. & Henikoff, S. Targeted in situ genome-wide profiling with high efficiency for low cell numbers. Nature protocols 13, 1006 (2018).

48. Cugusi, S., Kallappagoudar, S., Ling, H. & Lucchesi, J.C. The Drosophila helicase maleless (MLE) is implicated in functions distinct from its role in dosage compensation. Molecular & Cellular Proteomics 14, 1478–1488 (2015).

49. Herold, N. et al. Conservation of the protein composition and electron microscopy structure of Drosophila melanogaster and human spliceosomal complexes. Molecular and cellular biology 29, 281–301 (2009).

50. Lindehell, H., Kim, M. & Larsson, J. Proximity ligation assays of protein and RNA interactions in the male-specific lethal complex on Drosophila melanogaster polytene chromosomes. Chromosoma 124, 385–395 (2015).

51. Albig, C. et al. Factor cooperation for chromosome discrimination in Drosophila. Nucleic acids research 47, 1706–1724 (2019).

52. Quinn, J.J. et al. Revealing long noncoding RNA architecture and functions using domain-specific chromatin isolation by RNA purification. Nature biotechnology 32, 933–940 (2014).

53. Gabellini, N. A polymorphic GT repeat from the human cardiac Na+ Ca2+ exchanger intron 2 activates splicing. European Journal of Biochemistry 268, 1076–1083 (2001).

54. Hefferon, T.W., Groman, J.D., Yurk, C.E. & Cutting, G.R. A variable dinucleotide repeat in the CFTR gene contributes to phenotype diversity by forming RNA secondary structures that alter splicing. Proceedings of the National Academy of Sciences 101, 3504–3509 (2004).

55. Lin, C.-L. et al. RNA structure replaces the need for U2AF2 in splicing. Genome research 26, 12–23 (2016).

56. Hartmann, B. et al. Distinct regulatory programs establish widespread sex-specific alternative splicing in Drosophila melanogaster. Rna 17, 453–468 (2011).

57. Salz, H. & Erickson, J.W. Sex determination in Drosophila: The view from the top. Fly 4, 60–70 (2010).

58. Bell, L.R., Horabin, J.I., Schedl, P. & Cline, T.W. Positive autoregulation of sex-lethal by alternative splicing maintains the female determined state in Drosophila. Cell 65, 229–239 (1991).

59. Moschall, R. et al. Drosophila Sister-of-Sex-lethal reinforces a male-specific gene expression pattern by controlling Sex-lethal alternative splicing. Nucleic acids research 47, 2276–2288 (2019).

60. Haussmann, I.U. et al. m 6 A potentiates Sxl alternative pre-mRNA splicing for robust Drosophila sex determination. Nature 540, 301–304 (2016).

61. Moschall, R. (2019).

62. Horabin, J.I. & Schedl, P. Splicing of the drosophila Sex-lethal early transcripts involves exon skipping that is independent of Sex-lethal protein. Rna 2, 1–10 (1996).

63. Medenbach, J., Seiler, M. & Hentze, M.W. Translational control via protein-regulated upstream open reading frames. Cell 145, 902–913 (2011).

64. Wilkie, G.S., Dickson, K.S. & Gray, N.K. Regulation of mRNA translation by 5ʹ-and 3ʹ-UTR-binding factors. Trends in biochemical sciences 28, 182–188 (2003).

65. Petrova, V., et al. Chromatin accessibility regulates intron retention in a cell type-specific manner. bioRxiv (2021).

66. Agirre, E., Oldfield, A., Bellora, N., Segelle, A. & Luco, R. Splicing-associated chromatin signatures: a combinatorial and position-dependent role for histone marks in splicing definition. Nature communications 12, 1–16 (2021).

67. Newman, A.J. The role of U5 snRNP in pre-mRNA splicing. The EMBO journal 16, 5797–5800 (1997).

68. Graindorge, A., Carré, C. & Gebauer, F. Sex-lethal promotes nuclear retention of msl2 mRNA via interactions with the STAR protein HOW. Genes & development 27, 1421–1433 (2013).

69. Lin, J.-C. et al. The impact of the RBM4-initiated splicing cascade on modulating the carcinogenic signature of colorectal cancer cells. Scientific reports 7, 1–11 (2017).

70. Venables, J.P., Tazi, J. & Juge, F. Regulated functional alternative splicing in Drosophila. Nucleic acids research 40, 1–10 (2012).

71. Blencowe, B.J. Alternative splicing: new insights from global analyses. Cell 126, 37–47 (2006).

72. Mayne, B.T. et al. Large scale gene expression meta-analysis reveals tissue-specific, sex-biased gene expression in humans. Frontiers in genetics 7, 183 (2016).

73. Gibilisco, L., Zhou, Q., Mahajan, S. & Bachtrog, D. Alternative splicing within and between Drosophila species, sexes, tissues, and developmental stages. PLoS genetics 12, e1006464 (2016).

74. Revil, T., Gaffney, D., Dias, C., Majewski, J. & Jerome-Majewska, L.A. Alternative splicing is frequent during early embryonic development in mouse. BMC genomics 11, 399 (2010).

75. Aanes, H. et al. Differential transcript isoform usage pre-and post-zygotic genome activation in zebrafish. BMC genomics 14, 331 (2013).

76. Paris, M., Villalta, J.E., Eisen, M.B. & Lott, S.E. Sex bias and maternal contribution to gene expression divergence in Drosophila blastoderm embryos. PLoS Genet 11, e1005592 (2015).

77. Telonis-Scott, M., Kopp, A., Wayne, M.L., Nuzhdin, S.V. & McIntyre, L.M. Sex-specific splicing in Drosophila: widespread occurrence, tissue specificity and evolutionary conservation. Genetics 181, 421–434 (2009).

78. Lott, S.E., Villalta, J.E., Zhou, Q., Bachtrog, D. & Eisen, M.B. Sex-specific embryonic gene expression in species with newly evolved sex chromosomes. PLoS Genet 10, e1004159 (2014).

79. Ranz, J.M., Castillo-Davis, C.I., Meiklejohn, C.D. & Hartl, D.L. Sex-dependent gene expression and evolution of the Drosophila transcriptome. Science 300, 1742–1745 (2003).

80. Zhang, Y., Sturgill, D., Parisi, M., Kumar, S. & Oliver, B. Constraint and turnover in sex-biased gene expression in the genus Drosophila. Nature 450, 233–237 (2007).

81. Sun, X. et al. Sxl-Dependent, tra/tra2-Independent Alternative Splicing of the Drosophila melanogaster X-Linked Gene found in neurons. G3: Genes, Genomes, Genetics 5, 2865–2874 (2015).

82. Arbeitman, M.N., Fleming, A.A., Siegal, M.L., Null, B.H. & Baker, B.S. A genomic analysis of Drosophila somatic sexual differentiation and its regulation. Development 131, 2007–2021 (2004).

83. Bag, I., Dale, R.K., Palmer, C. & Lei, E.P. The zinc-finger protein CLAMP promotes gypsy chromatin insulator function in Drosophila. Journal of cell science 132 (2019).

84. Jordan, W. & Larschan, E. The zinc finger protein CLAMP promotes long-range chromatin interactions that mediate dosage compensation of the Drosophila male X-chromosome. bioRxiv (2020).

85. Vernes, S.C. Genome wide identification of Fruitless targets suggests a role in upregulating genes important for neural circuit formation. Scientific Reports 4, 4412 (2014).

86. Jai, Y.Y., Kanai, M.I., Demir, E., Jefferis, G.S. & Dickson, B.J. Cellular organization of the neural circuit that drives Drosophila courtship behavior. Current biology 20, 1602–1614 (2010).

87. Henninger, J.E. et al. RNA-mediated feedback control of transcriptional condensates. Cell 184, 207–225.e24 (2021).

88. Sharp, P.A., Chakraborty, A.K., Henninger, J.E. & Young, R.A. RNA in formation and regulation of transcriptional condensates. RNA 28, 52–57 (2022).

89. Oksuz, O. et al. Transcription factors interact with RNA to regulate genes. bioRxiv (2022).

90. Schneider, C.A., Rasband, W.S. & Eliceiri, K.W. NIH Image to ImageJ: 25 years of image analysis. Nature methods 9, 671–675 (2012).

91. Larschan, E. et al. Identification of chromatin-associated regulators of MSL complex targeting in Drosophila dosage compensation. PLoS Genet 8, e1002830 (2012).

92. Heyl, F., Maticzka, D., Uhl, M. & Backofen, R. Galaxy CLIP-Explorer: a web server for CLIP-Seq data analysis. GigaScience 9, giaa108 (2020).

93. DePristo, M.A. et al. A framework for variation discovery and genotyping using next-generation DNA sequencing data. Nature genetics 43, 491 (2011).

94. Andrews, S. FastQC: a quality control tool for high throughput sequence data. (Babraham Bioinformatics, Babraham Institute, Cambridge, United Kingdom, 2010).

95. Patro, R., Duggal, G., Love, M.I., Irizarry, R.A. & Kingsford, C. Salmon provides fast and bias-aware quantification of transcript expression. Nature methods 14, 417–419 (2017).

96. Yu, G., Wang, L.-G. & He, Q.-Y. ChIPseeker: an R/Bioconductor package for ChIP peak annotation, comparison and visualization. Bioinformatics 31, 2382–2383 (2015).

97. Ramírez, F., Dündar, F., Diehl, S., Grüning, B.A. & Manke, T. deepTools: a flexible platform for exploring deep-sequencing data. Nucleic acids research 42, W187–W191 (2014).

98. Bailey, T.L., Johnson, J., Grant, C.E. & Noble, W.S. The MEME suite. Nucleic acids research 43, W39–W49 (2015).

99. Langmead, B. & Salzberg, S.L. Fast gapped-read alignment with Bowtie 2. Nature methods 9, 357 (2012).

100. Kim, D., Paggi, J.M., Park, C., Bennett, C. & Salzberg, S.L. Graph-based genome alignment and genotyping with HISAT2 and HISAT-genotype. Nature biotechnology 37, 907–915 (2019).

101. Li, H. & Durbin, R. Fast and accurate short read alignment with Burrows–Wheeler transform. bioinformatics 25, 1754–1760 (2009).

102. Zhang, Y. et al. Model-based analysis of ChIP-Seq (MACS). Genome biology 9, 1–9 (2008).

103. Uyar, B. et al. RCAS: an RNA centric annotation system for transcriptome-wide regions of interest. Nucleic acids research 45, e91–e91 (2017).

104. Ge, S.X., Jung, D. & Yao, R. ShinyGO: a graphical gene-set enrichment tool for animals and plants. Bioinformatics 36, 2628–2629 (2020).

